# Molecular basis for the increased membrane fusion activity of the Ebola virus glycoprotein A82V variant from the 2013-2016 epidemic: insights from simulations and experiments

**DOI:** 10.1101/2024.10.28.620686

**Authors:** Natasha D. Durham, Aastha Jain, Angela Howard, Jeremy Luban, James B. Munro

## Abstract

During the 2013-2016 Ebola virus epidemic in Western Africa, an A82V mutation emerged in the envelope glycoprotein (GP) that persisted in most circulating isolates. Previous studies demonstrated that A82V increased GP-mediated membrane fusion and altered its dependence on host factors. The mechanistic basis for these observations, in particular the impact of A82V on the conformational changes in GP that are needed for membrane fusion, has not been evaluated in molecular detail. Here, using a combination of molecular dynamics simulations, fluorescence correlation spectroscopy, and a novel single-molecule Förster resonance energy transfer imaging assay, we specify the molecular mechanism by which A82V alters GP conformation to enhance GP-mediated viral entry. In so doing, we identify an allosteric network of interactions that links the receptor-binding site to the fusion loop of GP. Thus, the naturally occurring A82V mutation can tune the conformational dynamics of EBOV GP to enhance fusion loop mobility and subsequent viral fusion and infectivity in human cells.

## INTRODUCTION

The Ebola virus (EBOV) epidemic in Western Africa in 2013-2016 caused over 11,000 deaths, making it the largest documented filovirus outbreak. In early 2014, the A82V mutation arose in the EBOV envelope glycoprotein (GP), becoming a clade-defining variant that dominated subsequent circulating isolates throughout the outbreak. The A82V mutation showed increased pseudovirion tropism for a variety of human cell types *in vitro* (Diehl et al., 2016; Ueda et al., 2017; Urbanowicz et al., 2016). Further characterization of the mutation suggested increased functionality during interaction with endosomal membranes (Wang et al., 2017) and viral fusion (Diehl et al., 2016).

EBOV fusion naturally takes place in cellular endosomes. As a trimer of heterodimers, each GP protomer consists of a receptor-binding domain (GP1) and a membrane fusion domain (GP2) (Lee et al., 2008). GP1 contains the heavily glycosylated mucin-like domain and glycan cap, which conceal the binding site for the transmembrane protein and viral receptor Niemann-Pick C1 (NPC1) (Wang et al., 2016). Proteolysis by endosomal cathepsins remove the glycosylated domains, exposing the NPC1-binding site (Bornholdt et al., 2015; Chandran et al., 2005; Misasi et al., 2012) to generate a cleaved version of GP (GP^CL^). NPC1 binding alters interactions between GP1 and GP2 (Wang et al., 2016), which have been proposed to promote repositioning of GP1 with respect to GP2 to facilitate downstream fusion events (Durham et al., 2020). The acidic pH of the endosome facilitates conformational changes in GP that enhance the interaction of the fusion loop of GP2 with the endosomal membrane (Das et al., 2020; Gregory et al., 2014; Jain et al., 2023; Vaknin et al., 2024). Multiple studies also indicate a role for Ca^2+^ by enhancing GP interaction with target membranes and fusion *in vitro*, potentially by enabling engagement of the fusion loop with negatively charged phospholipids (Jain et al., 2023; Nathan et al., 2019; Odongo et al., 2023; Sakurai et al., 2015). At least one additional cleavage event by unidentified proteases is likely required for full membrane fusion (Fénéant et al., 2019; Markosyan et al., 2016; Odongo et al., 2023; Spence et al., 2016).

The mechanistic basis for the increased viral fusion and infectivity of the A82V mutation has been only partially resolved. Residue 82 is in the α1 helix of GP1, adjacent to the NPC1-binding site (Wang et al., 2016). Despite this proximity, no effect of A82V on NPC1 binding has been detected experimentally (Fels et al., 2021; Wang et al., 2017). Published observations support the hypothesis that A82V promotes conformational changes in GP that are relevant for membrane fusion. First, A82V increased resistance to the compound 3.47, which targets NPC1 and inhibits EBOV entry (Côté et al., 2011; Fels et al., 2021; Wang et al., 2017). This suggests that any NPC1-induced changes in GP structure may occur more readily in the presence of A82V. Second, the decreased melting temperature of A82V GP trimers compared to wild-type trimers suggests reduced structural stability overall (Fels et al., 2021; Wang et al., 2017). This relative instability was correlated with increased entry kinetics in live cells where fusion occurred earlier, in less acidic endosomes. This supports greater sensitivity to acidic pH, and thus more rapid pH-induced conformational changes with the A82V mutation. A structural rationale for this putative enhancement of fusion relevant conformational changes in GP remains elusive.

Here, we used molecular dynamics (MD) simulation of the GP ectodomain from the Makona strain of EBOV to generate hypotheses about the mechanistic basis for infectivity enhancement by A82V. Our simulations predict alterations in the NPC1-binding site stemming from the A82V mutation, which destabilizes the fusion loop through an allosteric network of interactions. Further analysis of essential dynamics indicated correlated motions consistent with increased fusion loop mobility for V82 GP. To experimentally corroborate the results of our MD simulations, we detected enhanced interaction of individual V82 GP trimers with target liposomal membranes. We also developed a novel single-molecule Förster resonance energy transfer (smFRET) imaging assay to probe EBOV pseudovirions for conformational changes in GP2 that reposition the fusion loop in response to conditions known to impact EBOV membrane fusion. Consistent with our hypothesis, the A82V mutation promoted large-scale conformational changes in GP2 at neutral pH that reposition the fusion loop. These data specify a model for EBOV infectivity enhancement by the A82V GP mutation at previously undefined molecular detail.

## RESULTS

### MD simulations assess the structural flexibility of wild-type (A82) and mutant (V82) GP^CL^ trimers

We first explored the structural basis for increased EBOV membrane fusion resulting from the A82V GP mutation in using MD simulation. We used molecular models of GP lacking the mucin-like domain and the glycan cap (GP^CL^) in these simulations since GP^CL^ is competent for NPC1 binding and downstream fusion-relevant conformational changes (Carette et al., 2011; Côté et al., 2011; Miller et al., 2012). Atomic coordinates for residues 32 to 188 of GP1 and 502 to 598 of GP2, taken from PDB 5JQ3, were used in MD simulations of the GP^CL^ trimer with either alanine at position 82 (A82) or valine at position 82 (V82). Each simulation was run in triplicate for 1μs each, yielding an aggregate sampling of 3μs for each protomer within the trimer. The difference in root mean square fluctuation (ΔRMSF) for each residue reflects the difference in flexibility of the A82 GP^CL^ trimer compared to the V82 trimer. Based on our ΔRMSF calculations, the A82V mutation increased the flexibility of the α1 helix (residues 79-85) and the upstream loop of GP1 (residues 70-78) (**Figure 1A-B**). This region contains residue 82 and flanks the NPC1-binding site. The increased flexibility of the α1 helix was due to the loss of helical content in the V82 simulation (26 ± 4%) as compared to A82 (44 ± 3%). A statistically significant (*p* < 10^−10^) decrease in helicity for this region was seen across all replicates (**Figure 1C-F**). The A82V mutation also increased dynamics throughout GP2, most notably within the fusion loop (residues 511-554) and the N-terminal portion of the HR1 helix (residues 555-565) (**Figure 1A-B**).

**Figure 1.**
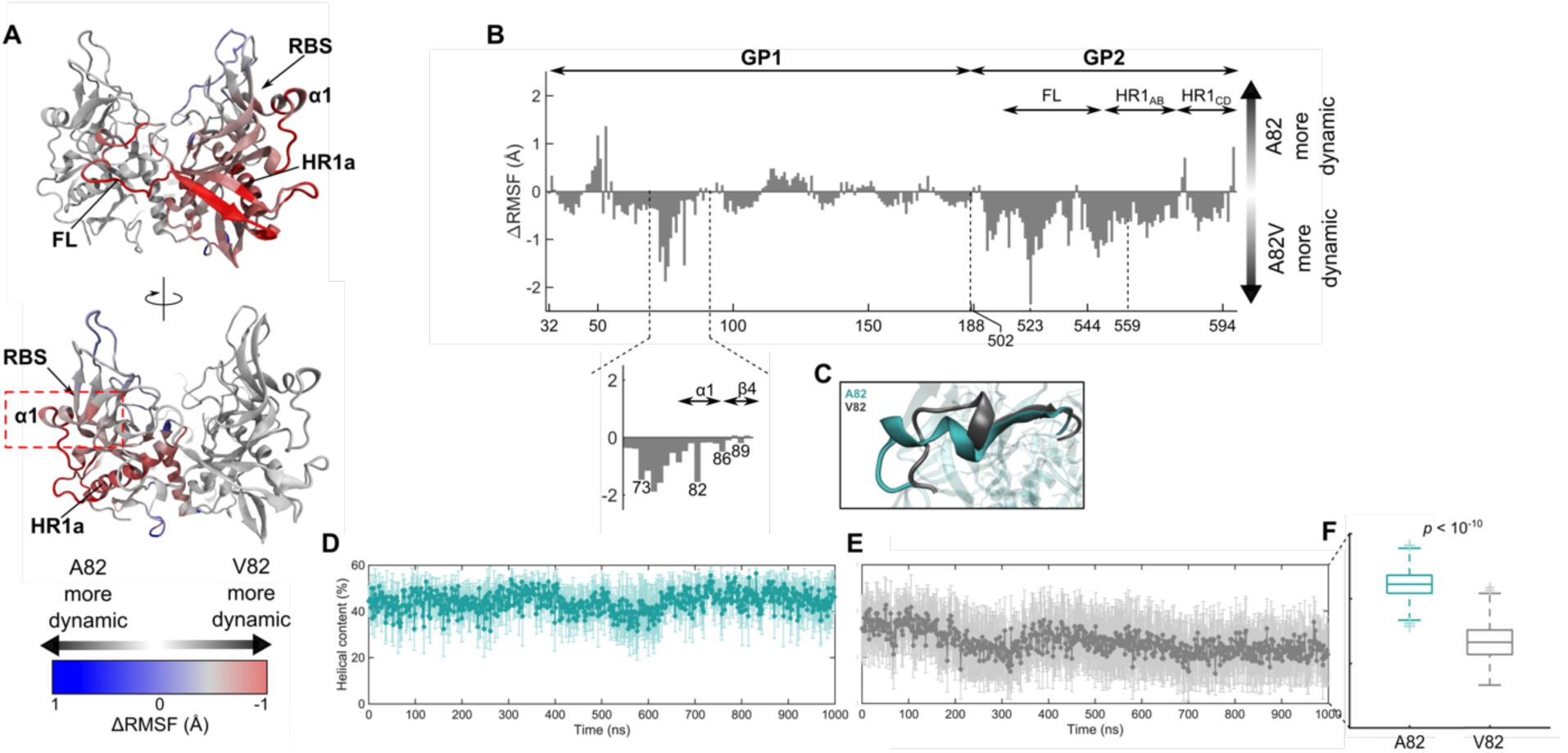
The A82V mutation increases the flexibility of GP in MD simulations. (**A**) The GP ectodomain structure with residues in one protomer colored according to the difference in RMSF between the A82 and V82 simulations (ΔRMSF = RMSF^A82^ – RMSF^V82^). Key structural elements are indicated: RBS, receptor-binding site; α1 helix; HR1, heptad repeat helix 1. (**B**) ΔRMSF for each residue in the GP ectodomain. The data are presented as the average of each protomer across the triplicate simulations, with error bars reflecting the standard error. (**C**) Visualization of the α1 helix in GP1 from representative frames in the A82 (cyan) and V82 (grey) simulations, indicating the loss of helical content in the V82 simulation. The region shown corresponds to the red dashed rectangle in panel (A). Quantification of the helical content of the α1 helix over the duration of the (**D**) A82 and (**E**) V82 simulations. Data are presented as the average of each protomer across the triplicate simulations, with error bars reflecting the standard error. (**F**) Box plots indicating the median, quantiles, and range of helical content seen across the simulations (A82, cyan; V82, grey). The *p*-value was determined by one-way ANOVA.

### Molecular interactions linking the GP1 receptor-binding site and GP2

We further analyzed the molecular interactions surrounding GP1 residues 79-85 of the α1 helix to investigate the origins of the decreased helicity in the V82 simulation. Compared to the A82 sidechain, the added bulk of the V82 sidechain led to reorientation of W86 and an unwinding of the α1 helix (**Figure 2**). The W86 sidechain sampled configurations within the receptor-binding site, including space that becomes occupied by F503 of NPC1 upon binding to GP^CL^ (**Figure 2E-F**) (Wang et al., 2016). Calculation of the sidechain dihedral angles of W86 in the V82 simulation indicated access to multiple configurations not seen for A82 (**Figure 2G-H**). The alternative configurations of W86 were visualized with a bivariate histogram displaying the distributions in W86 sidechain dihedrals and the distance between W86 and E178 (**Figure 2I**). The sidechain nitrogen of W86 engaged in stable electrostatic interactions with a sidechain oxygen of E178 in the A82 simulation. This interaction becomes less stable in the V82 simulation. The E178 sidechain also engaged in electrostatics with the H154 sidechain, a key sensor of endosomal pH (Jain et al., 2023). The backbone carbonyl of H154 in turn contacts the backbone amide nitrogen of Y534 within the fusion loop of the neighboring protomer in the trimer, thus providing a structural link between the receptor-binding site and the adjacent fusion loop. Consistent with this network of interactions, the motions of W86 were positively correlated with the motions of H154 and Y534 in the fusion loop of the neighboring protomer in both the A82 and V82 simulations (**Figure 3** and **Figure S1**). In summary, W86 is more mobile in the V82 simulation and gains access to multiple conformations. These dynamics translate to the fusion loop via interactions with H154, providing an allosteric link to the neighboring protomer.

**Figure 2.**
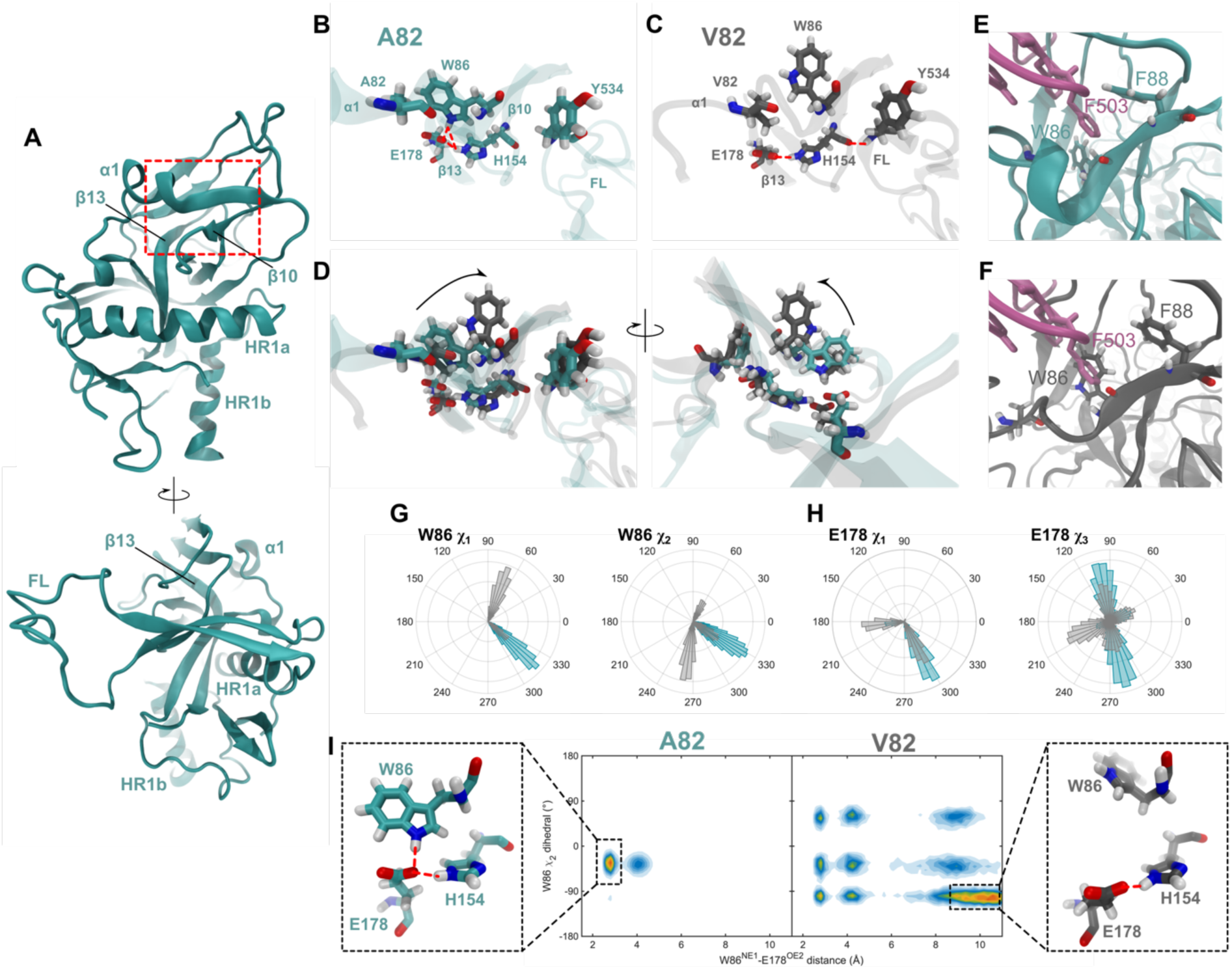
The A82V mutation reorients residue W86. (**A**) Structure of the GP^CL^ ectodomain monomer with key structural elements indicated (α1 helix; Δ10 and Δ13 sheets; HR1a and HR1b, the N- and C-terminal halves of the HR1 helix, respectively; FL, fusion loop). The red square indicates the region that is reflected in panels B through D. (**B**) Residues A82, W86 and nearby residues from a representative frame in the A82 simulation. Red dotted lines indicate putative electrostatic interactions between W86, H154, and E178. (**C**) A similar representative fame from the V82 simulation. (**D**) Overlays of the A82 (cyan), V82 (grey), and surrounding residues from two perspectives. The reorientation of the W86 side chain in the V82 simulation is indicated with a black arrow. (**E**) The receptor-binding site of GP with NPC1 (pink) bound indicating key hydrophobic residues (W86 and F88 in GP; F503 in NPC1) (PDB: 5F1B). (**F**) The same view of the A82V simulation with NPC1 docked into the equivalent position. (**G**) Polar histograms indicating the distribution of *X*_1_ and *X*_2_ sidechain dihedral angles of W86 in the A82 (cyan) and A82V (grey) simulations. (**H**) Polar histograms indicating the distribution of *X*_1_ and *X*_3_ sidechain dihedral angles of E178 in the A82 (cyan) and V82 (grey) simulations. (**I**) Bivariate histograms indicating the existence of multiple conformations of the residues highlighted in panels B through D. Structural depictions of the predominant conformations are indicated for the A82 and V82 simulations.

**Figure 3.**
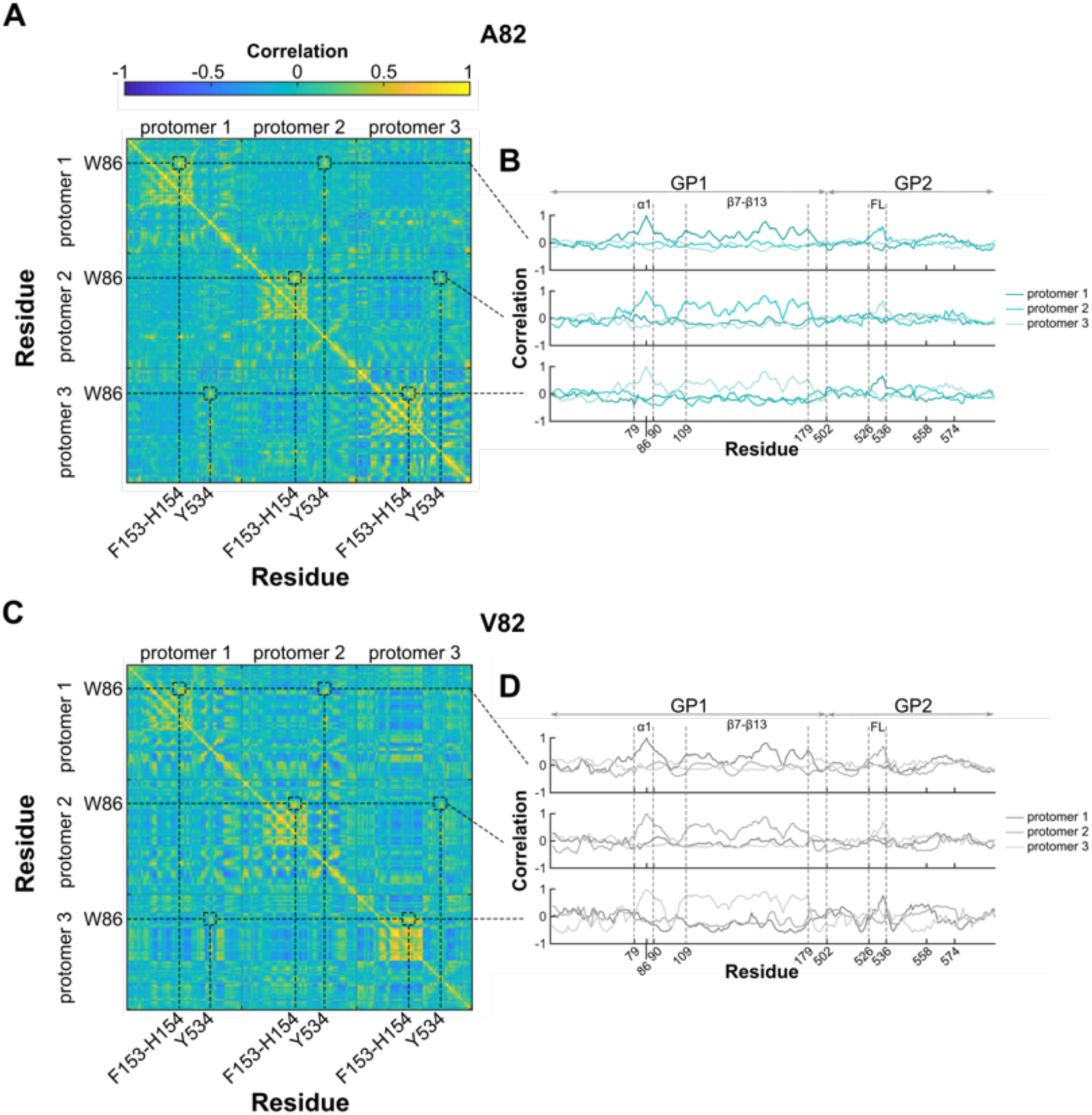
Correlation of W86 and GP^CL^ fusion loop dynamics. (**A**) Correlation matrix of all residues in the GP^CL^ A82 trimer determined from one simulation replicate. The colorbar indicates the correlation coefficient. Key residues within the three protomers of the GP^CL^ trimer are highlighted. (**B**) Correlation coefficient of W86 position with all residues within each of the three protomers. Key residues and structural features are indicated. High correlation is seen between W86 and the α1 helix (cc ≈ 0.8) and Δ7-Δ13 sheets (including H154 in Δ10; cc > 0.8) in the same protomer, and the fusion loop (FL) of a neighboring protomer (cc ≈ 0.6). (**C**) Correlation matrix of all residues in the GP^CL^ V82 trimer, displayed as in (A). (**D**) The correlation coefficient of W86 position with all other residues in the GP^CL^ V82 trimer, displayed as in (B). Equivalently high correlation was seen between W86 and the α1 helix, Δ7-Δ13 sheets, and the FL as for the A82 simulations. The same analysis from the additional two replicates of each simulation is shown in **Figure S1**.

The increased flexibility of the α1 helix in the V82 simulation also influenced motions of upstream residues. In A82 GP^CL^, residue N73 in the 3^10^ helix of the Δ3-α1 loop forms stable electrostatic contacts with the K510 backbone and the R559 sidechain, which reside N-terminal to the fusion loop and in the HR1 helix of GP2, respectively (**Figure 4 A-B**). These interactions appear to stabilize the fusion loop and HR1 prior to fusion triggering (Bornholdt et al., 2015; Lee et al., 2008; Wang et al., 2016; Zhao et al., 2016). For A82 GP^CL^, we observed a predominant orientation of the N73 sidechain with minor sampling of alternative configurations. The loss of helicity of the α1 helix in the V82 simulation increased the mobility of N73, destabilizing its interactions with K510 and R559. As a result, multiple orientations of the N73 sidechain with respect to K510 and R559 were seen (**Figure 4C-F**). Taken together, these data predict that the A82V mutation destabilizes the fusion loop in its hydrophobic cleft.

**Figure 4.**
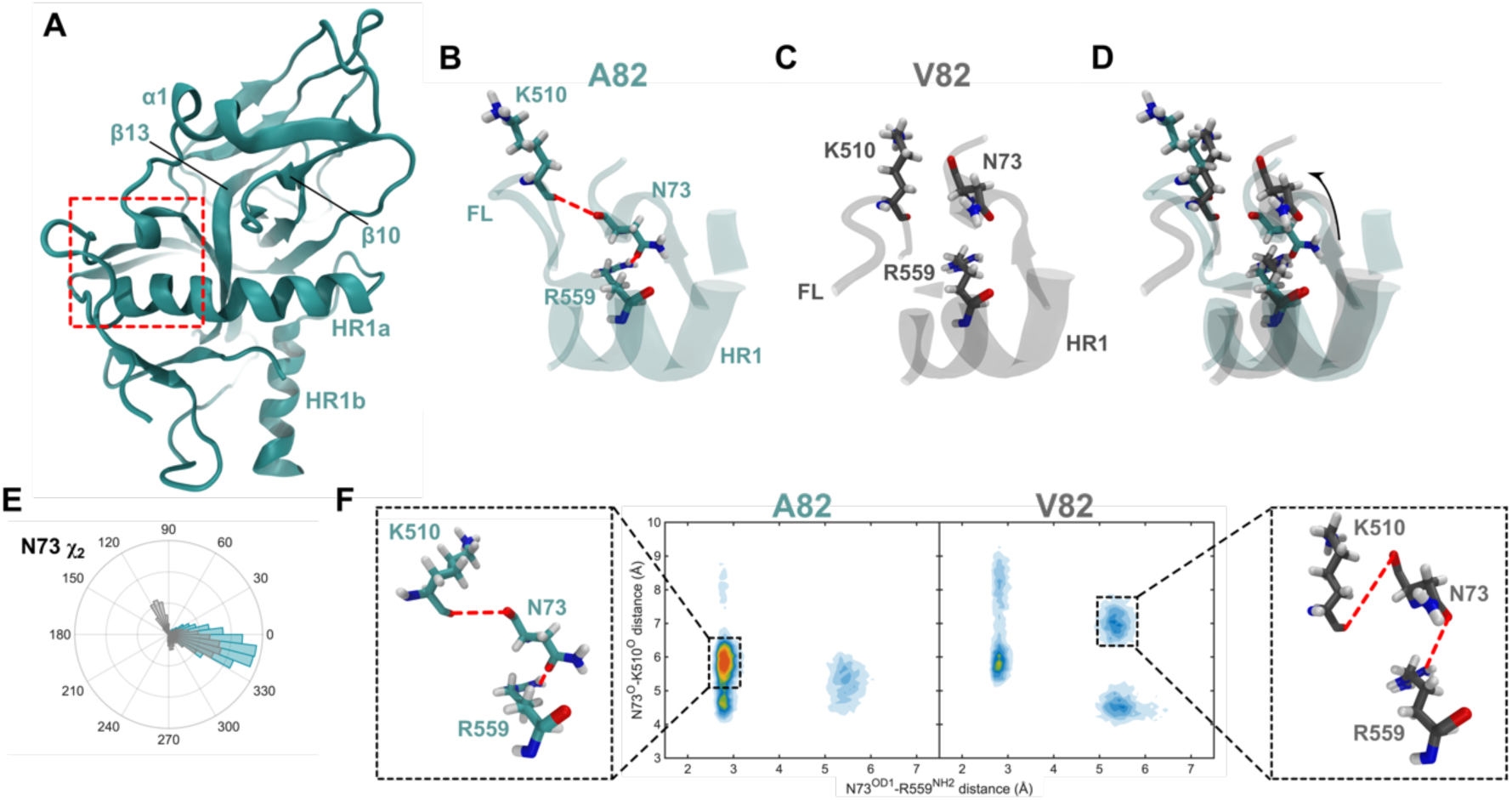
A82V disengages N73 from GP2. (**A**) GP^CL^ ectodomain monomer with key structural elements indicated (α1 helix; Δ10 and Δ13 sheets; HR1a and HR1b, the N- and C-terminal halves of the HR1 helix, respectively). The red square indicates the region that is reflected in panels B through D. (**B**) Zoomed-in view of N73, K510, and R559 from a representative frame in the A82 simulation. Red dotted lines indicate putative electrostatic interactions between N73 and K510, and with R559. (**C**) The same zoomed-in view from the V82 simulation. (**D**) Two perspectives of the overlay from the A82 (cyan) and V82 (grey) simulation. The reorientation of the NM73 side chain in the A82V simulation is indicated with a black arrow. (**E**) Polar histogram indicating the distribution of the *X*_2_ sidechain dihedral angle of N73 in the A82 (cyan) and V82 (grey) simulations. (**F**) Bivariate histograms indicating the existence of multiple conformations of the residues highlighted in panels B through D. Structural depictions of the predominant conformations are indicated for each mutant.

### The A82V mutation increases mobility of the GP2 fusion loop

To address whether the destabilizing interactions above led to greater movement of the fusion loop in the V82 simulation, we used principal component analysis to extract the collective motions of the GP^CL^ trimer from the simulations (Amadei et al., 1993). The first three principal components reflected approximately 40% of the total variance in atomic positions seen in the simulations (**Figure S2**). To identify GP^CL^ trimer conformations sampled in the simulations, each simulation frame was projected into the space formed by the first three principal components (**Figure 5A-C**). The frames were then clustered using a Gaussian mixture model yielding 9 GP^CL^ conformations for each of the two simulations. The RMSDs of the fusion loops for each of the 9 conformations was greater for V82 than for A82 (**Figure 5D**). For each conformation, we also determined the distances between the centers of mass of the three fusion loops in the trimer. In the symmetric GP^CL^ structure that served as the starting point of the simulations, this distance was 42 Å. In the A82 simulations, the top two most populated conformations, which together comprise 44% of the sampling, indicated fusion loop movements toward the central trimer axis, which reduced distances between the fusion loops (**Figure 5E**). In contrast, the two most populated V82 conformations, which comprise 49% of the sampling, indicated outward movement of the fusion loops away from the central trimer (**Figure 5F**). Overall, the V82 simulation showed greater movement of the fusion loops away from the central trimer axis compared to A82.

**Figure 5.**
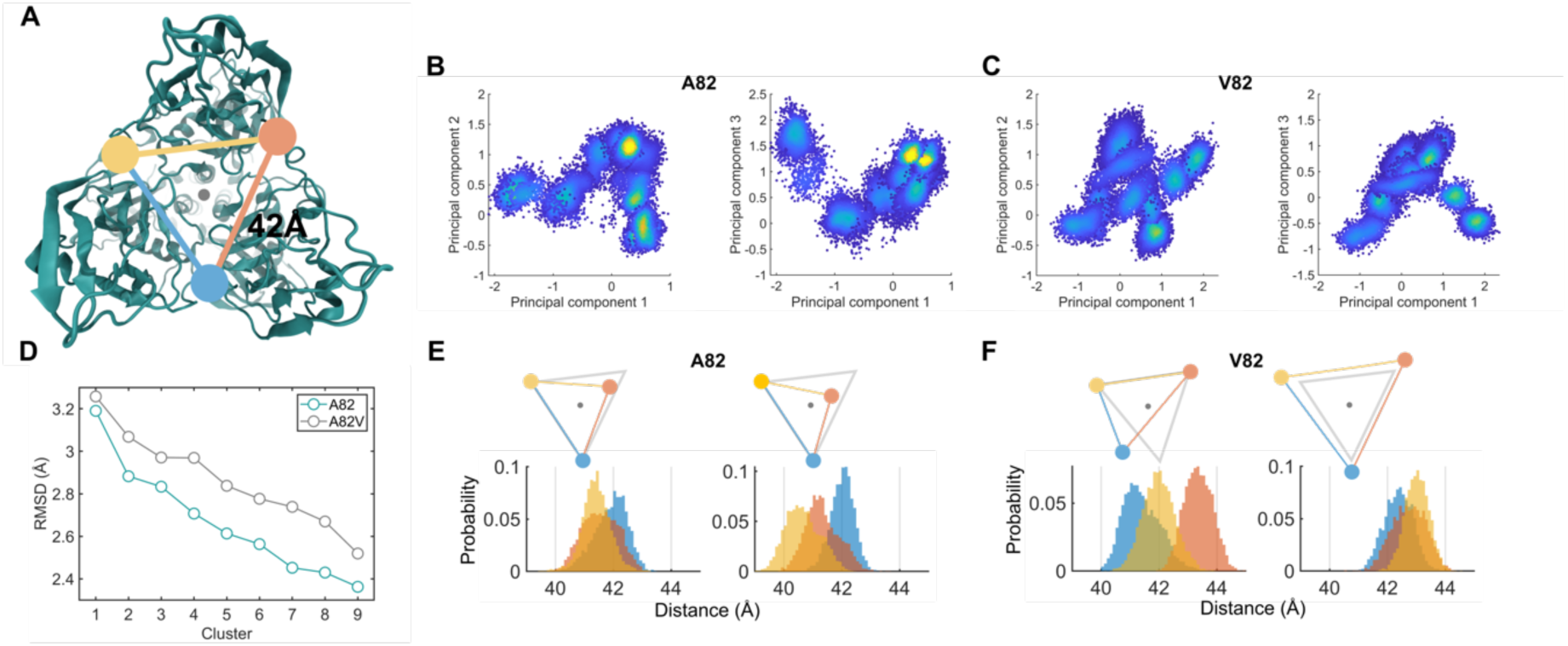
Principal component analysis indicates distinct conformations of A82 and V82 GP^CL^ trimers. (**A**) The symmetric GP^CL^ trimer, which served as the starting points for the MD simulations, with the centers of mass of the three fusion loops indicated by colored circles. The distance between the three fusion loops is indicated (42 Å). (**B**) Projection of the frames of the concatenated MD simulations (approximately 18,000 frames) of GP^CL^ A82 into the space formed by the first three principal components. Clustering of the frames identified nine conformations sampled in the simulations. Points are colored according to their density in each cluster with blue and yellow indicating low and high density, respectively. (**C**) The same data for V82 GP^CL^. (**D**) The RMSD of each of the nine conformations identified for A82 (cyan) and V82 (grey). (**E**) The distribution of distances between the centers of mass of the three fusion loops in each of the three protomers in the A82 GP^CL^ trimer, colored according to the triangle in (A). The distance distributions for the two most populated clusters are shown. Also shown for each set of distributions is a triangular diagram indicating the approximate movement of the fusion loops as compared to symmetric starting point (grey). (**F**) The same analysis for V82 GP^CL^.

### The A82V mutation increases GP^CL^ interaction with membranes

We next asked if movement of the fusion loop away from the trimer axis could position the V82 GP^CL^ fusion loop to engage with a target membrane more readily. To address this question experimentally, we used an established fluorescence correlation spectroscopy (FCS) assay to probe the interaction between the trimeric GP^CL^ ectodomain (GP^CL^ΔTM) and model membranes. We previously demonstrated that this assay reports on pH- and Ca^2+^-dependent GP fusion loop-mediated interaction with liposomes containing anionic phospholipids characteristic of the late endosome (**Figure 6A**, Materials and Methods) (Jain et al., 2023). At neutral pH, the A82V mutation increased the fraction of membrane-bound GP^CL^ΔTM by approximately 5%, which further increased to nearly 10% in the presence of 1 mM Ca^2+^ (**Figure 6B**). No significant effect of the A82V mutation on the GP-membrane interaction was seen at pH 5.5, irrespective of the Ca^2+^ concentration.

**Figure 6.**
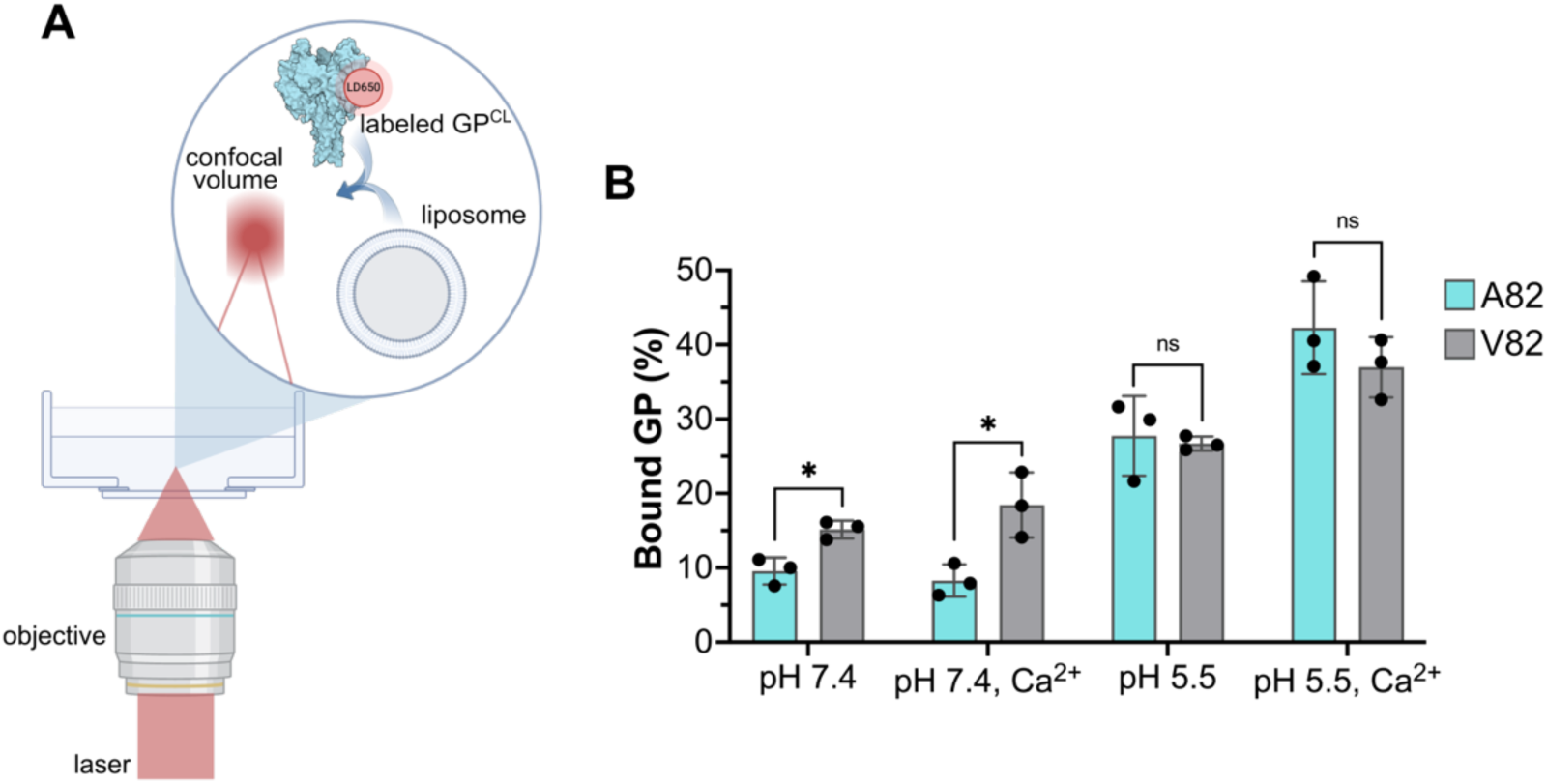
GP^CL^ trimer-membrane interaction by FCS. (**A**) Schematic of the established FCS assay for probing the interaction between fluorescently labeled GP and a liposome (Jain et al., 2023). (**B**) The fraction of GP^CL^ bound to the liposomes under the indicated conditions for A82 (cyan) and V82 (grey). Bars indicate the mean of three technical replicates with error bars indicating the standard error. Asterisks indicate *p* < 0.05, determined by *t* test (ns, *p* > 0.05).

### smFRET imaging reveals conformational changes in GP2

Based on our MD, which indicate greater mobility of the fusion loop in the presence of the A82V mutation, and our FCS experiments, which indicate greater association between the V82 GPΔTM and the target membrane, we further hypothesized that this mutation could lead to conformational changes in GP2. To test this hypothesis, we developed a smFRET imaging assay to report on the conformational changes that reposition the fusion loop. We generated pseudoparticles containing GP^CL^ with donor and acceptor fluorophores attached to a single fluorescently labeled protomer at amino acid positions 550 and 617, which are located within the fusion loop and at the N-terminal edge of the HR2 helix, respectively. Pseudoparticles displaying GP^CL^ were immobilized on a quartz microscope slide and imaged using total internal reflection fluorescence (TIRF) microscopy (**Figure 7A-B**). We initially collected smFRET trajectories from individual A82 GP^CL^ protomers on the pseudovirion surface at pH 7.5. To determine the number of FRET states observed in the trajectories, we fit the trajectories to several hidden Markov models (HMMs) using maximum likelihood estimation (Qin et al., 2000) and compared the model fitness using the Akaike information criterium (AIC; **Figure S3**) (Akaike, 1974). The model that best represented the smFRET trajectories contained 4 non-zero FRET states with means and standard deviations of 0.8±0.1, 0.5±0.1, 0.12±0.1, and 0.05±0.06. The model also contained a 0-FRET state that accounts for photobleaching and conformations where the distance between the fluorophores is beyond the detectable range. The 0.8-FRET state had the greatest occupancy (53±2%; **Figure 7C-D**, **Table 1**). MD simulation of labeled GP^CL^ in the prefusion conformation indicated an average distance between the fluorophores of 36±5 Å, which is consistent with the predominant 0.8-FRET state (**Figure S4**) (Bornholdt et al., 2015; Lee et al., 2008; Zhao et al., 2016). The observed transitions out of the 0.8-FRET predominant prefusion state to lower FRET states suggest alternative conformations in which the fusion loop has moved out of the hydrophobic cleft, increasing the distance between the fluorophores. To quantify the conformational fluctuations of GP^CL^, we used the HMM analysis of the smFRET trajectories to determine the rates of transition out of the 0.8-FRET prefusion conformation. We extracted the dwell times in the 0.8-FRET pre-fusion conformation from the results of the HMM analysis of GP^CL^ with A82 and A82V, and compiled them into histograms. For A82 at neutral pH, the histograms were fit to the sum of two exponential functions (*A*_fast_ exp (*t*/*t*_fast_) + *A*_slow_ exp (*t*/*t*_slow_)) with time constants *t*_fast_ = 0.7±0.2 s and *t*_slow_ = 4±1 s with amplitudes of *A*_fast_ = 59±5% and *A*_slow_ = 41±5%, respectively (**Figure 8A**). The double exponential dwell time histogram indicates the existence of two pathways for transition out of the pre-fusion conformation.

**Figure 7.**
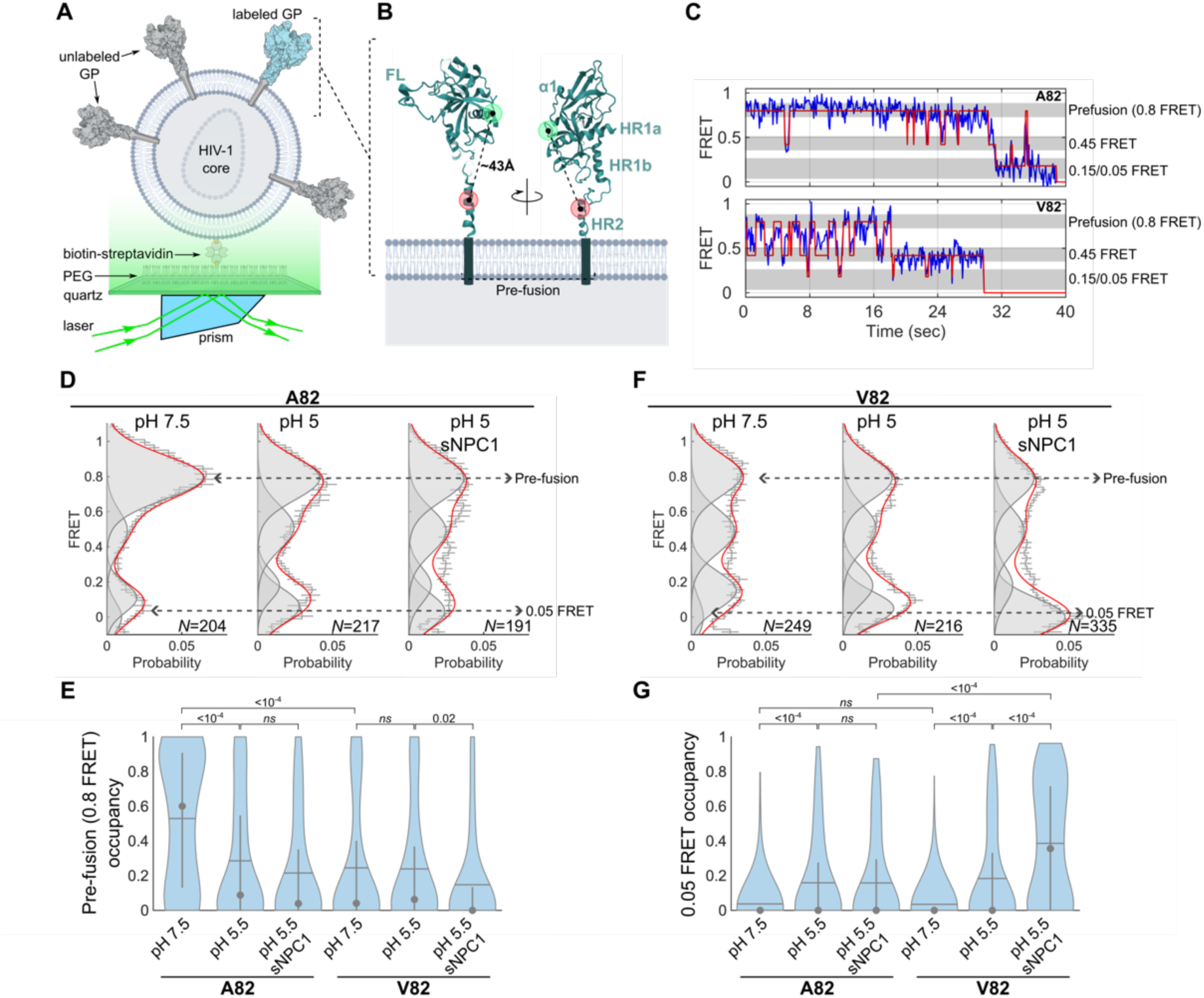
smFRET imaging reveals GP^CL^ conformations stabilized and destabilized by A82V. (**A**) Schematic of the smFRET assay in which pseudovirions formed with the HIV-1 core are immobilized to enable imaging with TIRF microscopy. (**B**) A GP^CL^ monomer with the sites of fluorophore attachment indicated (donor, green; acceptor, red. PDB: 5JQ3). (**C**) Example smFRET trajectories from individual GP^CL^ trimers with (top) A82 or (bottom) V82. The experimental trajectories are shown in blue, overlaid with the idealized trajectory resulting from fitting to the 5-state HMM (red). The prefusion conformation is indicated with a grey bar, along with the observed intermediate conformations. (**D**) FRET histograms for A82 GP^CL^ smFRET trajectories (N)were acquired under the indicated conditions. Histograms reflect the average of three independent groups of trajectories; error bars represent the standard error. Overlaid on the histograms are five Gaussian distributions (grey) with means determined through HMM analysis of the individual smFRET trajectories, along with the sum of the Gaussians shown in red. The 0.8-FRET pre-fusion conformation and the 0.05-FRET state are indicated. (**F**) FRET histograms for V82 GP^CL^ as in (D). (**E**) Violin plots displaying the distribution in occupancy in the 0.8-FRET pre-fusion conformation in each GP^CL^ population imaged under the indicated condition. Horizontal lines indicate the population mean occupancies; the grey circles and whiskers indicate the medians and quantiles, respectively. *p*-values are indicated, which were determined by 1-way ANOVA and multiple comparison testing (ns, *p* > 0.05). (**G**) Violin plots displaying the distribution in occupancy in the 0.05-FRET state, displayed as in (E). Numeric data are presented in **Table 1**.

**Figure 8.**
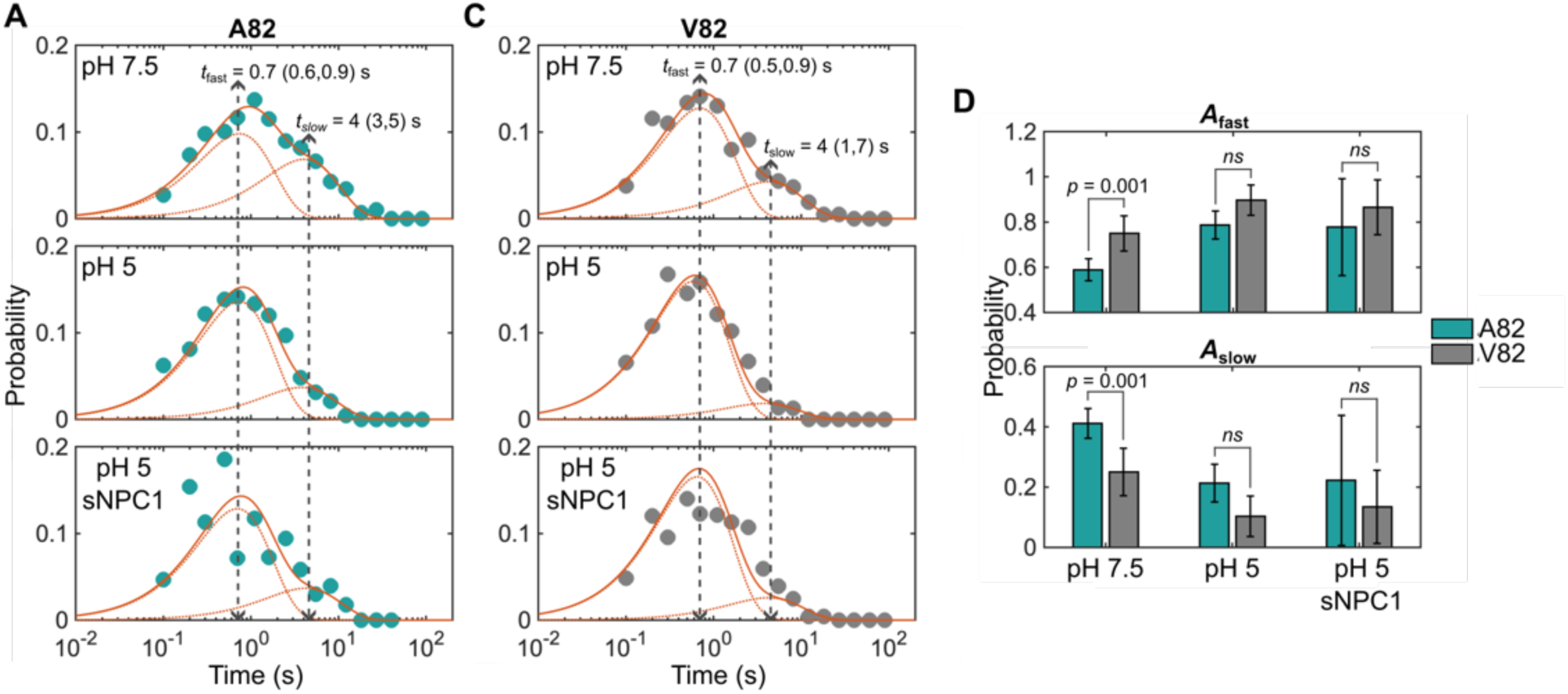
Kinetic analysis of the conformational transitions determined by smFRET. Histograms of dwell times in the 0.8-FRET pre-fusion conformation extracted from the HMM analysis of smFRET trajectories (blue circles) acquired for (**A**) A82 and (**C**) V82 GP^CL^, under the indicated conditions. Histograms are displayed with logarithmically spaced bins to assist in visualization of the two time constants (Sigworth and Sine, 1987). Histograms were fit to double exponential functions *A*_fast_ exp (*t*/*t*_fast_) + *A*_slow_ exp (*t*/*t*_slow_), where *A*_fast_ and *A*_slow_ are amplitudes, and *t*_fast_ and *t*_slow_ are the corresponding time constants. Double exponential fits are overlaid in solid red lines with the individual exponentials in dashed red lines. The two time constants are indicated with 95% confidence intervals in parentheses. (**D**) The amplitudes, *A*_fast_ and *A*_slow_, determined in the exponential fitting in (A) and (B). Bars reflect the fitted values with error bars indicating the 95% confidence intervals. *p*-values are indicated (ns, *p* > 0.05).

**Table 1.**
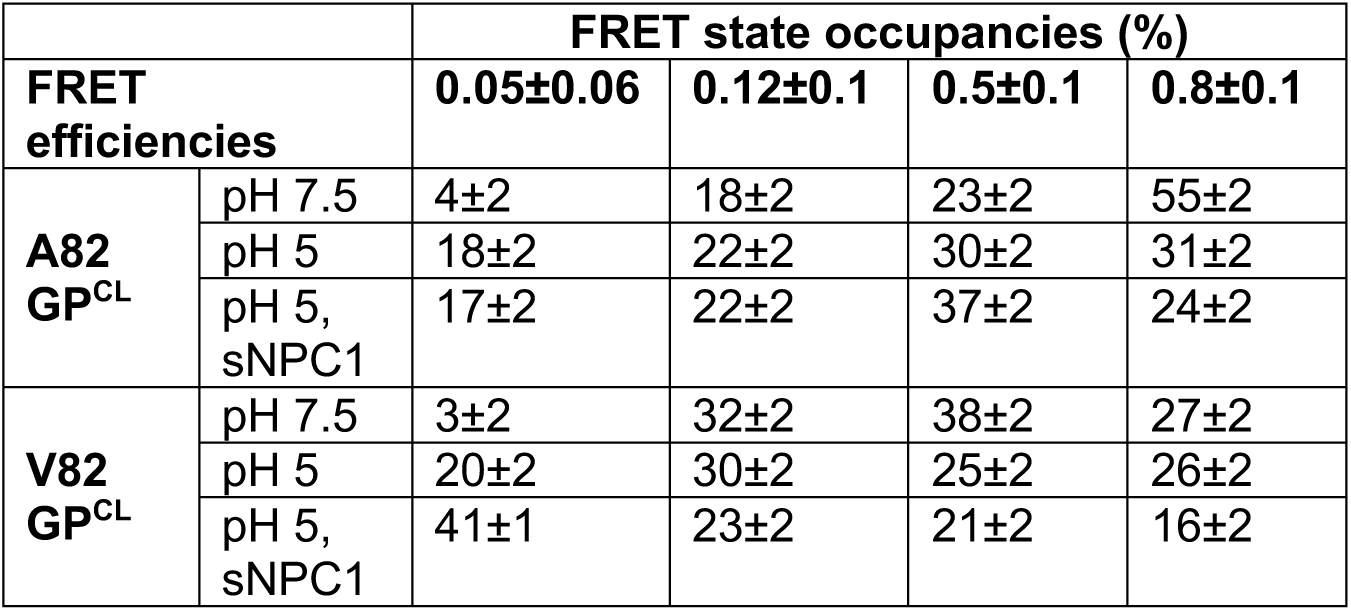
FRET state occupancies determined through HMM analysis. FRET efficiencies are shown as the mean ± standard deviation. Occupancies are presented as the mean ± SEM.

We next determined the impact of acidic pH on GP2 conformational changes. Imaging at pH 5.5 decreased the occupancy of the 0.8-FRET state (28±2%) and increased the occupancy in the 0.05-FRET state (16±2%). We observed the same time constants, *t*_fast_ and *t*_slow_, as seen at neutral pH but with amplitudes shifted to *A*_fast_ = 79±6% and *A*_slow_ = 21±6%. Thus, acidic pH promoted the fast kinetic pathway for transition out of the pre-fusion conformation. The addition of the soluble domain C of NPC1 (sNPC1) had a slight effect on the FRET state occupancies but was not statistically significant (**Figure 7D-E**, **Table 1**). Likewise, we detected no effect on the kinetics of transition out of the 0.8-FRET state. Therefore, consistent with our previous reports (Das et al., 2020; Jain et al., 2023), these data suggest that the GP^CL^ fusion loop is repositioned by conformational changes induced by acidic pH.

### The A82V mutation promotes conformational changes in GP2

We next asked whether the A82V mutation affected the conformational dynamics of GP2, as predicted by our MD simulations and membrane-binding data. smFRET data acquired for GP^CL^ with A82V at pH 7.5 indicated the same FRET states that were present for A82 GP^CL^. However, the A82V mutation significantly reduced the occupancy in the 0.8-FRET pre-fusion state to 24±2%, compared to 53±2% for A82 (*p* < 10^−4^). Analysis of the kinetics of transitions out of the predominant prefusion conformation again indicated the same time constants, *t*_fast_ and *t*_slow_, as seen for A82. However, the amplitudes were shifted toward the fast time constant with *A*_fast_ = 75±8% and *A*_slow_ = 25±8% (**Figure 8C**). These amplitudes were significantly different from the amplitudes seen for A82 at neutral pH (*p* = 0.001; **Figure 8D**). This indicates that the A82V mutation promotes the fast kinetic pathway out of the pre-fusion conformation associated with the 0.8 FRET state.

Acidic pH did not further affect the occupancy of the 0.8-FRET state of V82 GP^CL^. However, the occupancy in the 0.05-FRET state was increased to 20±2%. Addition of sNPC1 to V82 GP^CL^ further reduced the occupancy of the pre-fusion conformation, although the change in kinetics failed to reach statistical significance. A more pronounced impact of sNPC1 binding was seen on the occupancy in the 0.05-FRET state, which increased to 39±1%. The occupancy in the 0.05-FRET state may be under-represented due to our limited ability to differentiate it from 0 FRET. Thus, sNPC1 binding had a greater impact on the conformation of V82 GP^CL^ than on A82 GP^CL^. Taken together, these data demonstrate that at neutral pH, the conformational equilibrium of the V82 GP^CL^ trimer is distinct from the A82 GP^CL^ trimer. Furthermore, the A82V mutation makes GP^CL^ less reliant on acidic pH to induce conformational changes and more responsive to the conformational effects of sNPC1-C binding.

## DISCUSSION

Our results indicate that the A82V GP mutation enhances EBOV infectivity by destabilizing the predominant prefusion conformation of GP^CL^ and promoting conformational changes that are on pathway to membrane fusion. This conclusion corroborates other reports of decreased thermal stability of GP^CL^ in the presence of the A82V mutation (Fels et al., 2021; Wang et al., 2017). Our MD simulations provided insights into the molecular basis for this destabilization of prefusion GP^CL^. The simulations indicated that V82 alters the secondary structure of the α1 helix in GP1, and these structural alterations were propagated through electrostatic interactions with H154 to Y534 in the fusion loop of the neighboring protomer. The presence of V82 also affected the HR1 helix of GP2, destabilizing inter-domain interactions, resulting in greater fusion loop dynamics and propensity to move outward, away from the trimer axis. We also observed increased engagement with a target membrane, which is supported by a previous report (Wang et al., 2017). This increased GP^CL^-membrane interaction may result from the greater mobility of the fusion loop. Finally, our smFRET imaging data provide direct observation of conformational changes promoted by the A82V mutation. These data demonstrate that the residue V82 promotes movement of the fusion loop in GP^CL^, similar to the effect of acidic pH on A82 GP^CL^.

Analysis of the A82V mutation has reinforced the importance of an allosteric network of electrostatic interactions that links the receptor-binding site to the fusion loop. Our previous study demonstrated that residue H154 in GP1 senses pH changes relevant to viral trafficking to the late endosome (Jain et al., 2023). We reported that protonation of H154 led to destabilization of electrostatic interactions with Y534 in the fusion loop. The present study of A82V suggests that the same allosteric network is utilized for promotion of conformational changes that reposition the fusion loop. That is, the A82V mutation plays a similar functional role as protonation of H154. Our FCS and smFRET results demonstrate that V82 GP^CL^ has a conformational equilibrium at neutral pH that resembled the equilibrium of A82 GP^CL^ at acidic pH, implying V82 GP^CL^ has decreased dependence on acidic pH to promote conformational changes necessary for fusion. Previous live-cell imaging experiments demonstrated that pseudovirions with the A82V GP mutation had decreased the time to fusion, suggesting entry through an earlier, less acidic endosome (Fels et al., 2021). Taken together, these results suggest that the A82V GP mutation enables EBOV fusion to occur under less acidic conditions.

Previous investigations have not detected any direct effect of the V82 residue on GP^CL^ binding to NPC1. However, A82V reduced the dependence of GP^CL^ on NPC1 for fusion triggering during virus entry (Wang et al., 2017). This conclusion was based largely on reduced sensitivity to 3.47, an inhibitor that antagonizes the GP^CL^-NPC1 interaction (Côté et al., 2011). The localized changes in GP^CL^ conformation seen in our MD simulations may facilitate, or possibly emulate, the structural alterations following NPC1 binding. The structure of the sNPC1-bound wild-type GP^CL^ shows an upward shift in the α1 helix as compared to unbound GP^CL^ (Wang et al., 2016). The α1 helix movement may assist with stabilization of the GP1-NPC1 interface. In particular, the α1 helix shift coincides with reorientation of T83, which forms stabilizing electrostatic interactions with D129 of NPC1. The increased flexibility of the α1 helix seen in the V82 simulation may facilitate its displacement during NPC1 binding such that these stabilizing interactions form more readily. The sNPC1-bound GP^CL^ structure also indicates reorientation of N73 and loss of electrostatic interactions with K510 and R559. This led to an apparent repulsion between K510 and R559, which may contribute to releasing the fusion loop from the hydrophobic cleft. While our V82 GP^CL^ simulation does not indicate repulsive interactions between K510 and R559, the added flexibility resulting from disengagement of N73 could facilitate K510 adopting an orientation that creates repulsion with R559 following NPC1 binding. The loss of the stabilizing contact between N73 and K510 may on its own contribute to destabilization of the fusion loop.

Kinetic analysis of the transition rates out of the pre-fusion conformation were not changed by the A82V mutation. Both A82 and V82 GP^CL^ underwent transitions out of the predominant 0.8 FRET prefusion conformation with the same time constants, *t*_fast_ and *t*_slow_. The destabilization of this prefusion conformation induced by A82V was due to greater amplitude of the fast transitions. The consistency in the time constants for transition out of the prefusion conformation indicates that the energetics of this transition are the same for both A82 and V82, which suggests that the structural nature of the conformational change is comparable. In other words, V82 does not promote a GP^CL^ conformational change that is inaccessible to A82. Rather V82 promotes an accelerated kinetic pathway to exposure of the fusion loop and engagement with the target membrane. The structural distinctions of the *fast* and *slow* transitions are not known but may involve the inter-domain and inter-protomer interactions implicated by our MD simulations.

This study demonstrates how quantitative biophysical interrogations can reveal the mechanistic underpinnings of mutations that enhance viral infectivity. Future studies of mutations that evolve in response to interspecies transmission, changing immune status, or the introduction of antivirals will benefit from similar investigations.

## ACKNOWLEDGEMENTS

This work was supported by National Institutes of Health grants R01AI174645 (to J.B.M.) and R01AI148784 (to J.L.).

## AUTHOR CONTRIBUTIONS

N.D.D. and J.B.M.: conceptualization; N.D.D., A.J., and J.B.M.: methodology; N.D.D., A.J., and J.B.M.: formal analysis; N.D.D., A.J., A.H., and J.B.M.: investigation; J.L.: resources; N.D.D. and J.B.M.: data curation; N.D.D. and J.B.M.: writing – original draft; N.D.D., A.J., J.L., and J.B.M.: writing – review & editing; N.D.D. and J.B.M.: supervision; J.L. and J.B.M.: funding acquisition.

## STAR METHODS

### MD simulations of GP^CL^ with A82 and V82

An atomic model of trimeric GP^CL^ with A82, including resides 32 to 188 of GP1 and 502 to 598 of GP2, was generated using coordinates from PDB 5JQ3 (Zhao et al., 2016). A second GP^CL^ model was generated containing the A82V mutation using the Mutator plugin in VMD. In both cases, the N and C termini of GP1 were capped with neutral ACE and NME moieties. The C terminus of GP2 was similarly capped with NME. Missing atoms and explicit solvent, including Na^+^ and Cl^−^ ions to a final concentration of 150 mM, were added using the LEaP program in AmberTools. The complete system was comprised of approximately 77,000 atoms. The protein and solvent components of the model were parameterized in LEaP using the Amber (ff19SB) and the TIP3P forcefields, respectively. Following 10,000 steps of conjugate gradient energy minimization, the systems were equilibrated through multiple 0.3-ns simulations with a 1 fs time step. First, the protein backbone atoms were fixed and run with the temperature increasing from 100K to 300K in 100K increments. Next, harmonic restraints were imposed on the backbones and released gradually over four simulations at 300K with force constants 1, 0.5, 0.1, and 0.01 kcal/(mol Å^2^). Finally, the systems were then simulated without restraints for an additional 10 ns with a 2 fs time step. The production simulations were run in the NPT ensemble with rigid bonds enforced and periodic boundary conditions. The temperature was controlled at 300K with the Langevin thermostat. Pressure was maintained with the Nose-Hoover Langevin piston at 1 atm. Electrostatic interactions were determined using the particle mesh Ewald method and van der Waals interactions were calculated with a cutoff distance of 10 Å. Simulations were run in triplicate for 1 μs each with a GPU-accelerated installation of NAMD v3.0b2 on a GPU node on the SCI cluster at UMass Chan Medical School.

### MD simulations of dye mobility

To estimate the space sampled by the fluorophores, an atomic model of trimeric A82 GP^CL^ was generated from PDB 5JQ3, which included residues 32 to 188 of GP1 and 502 to 631 of GP2. Cy3 and Cy5 fluorophores were attached at positions 550 and 617 in CHARMM-GUI. As above, N and C termini were capped with ACE and NME moieties, and missing atoms and explicit solvent, including Na^+^ and Cl^−^ ions to a final concentration of 150 mM, were added. The protein and the fluorophores were parameterized with the CHARMM36m and CGenFF forcefields, respectively. And the solvent was parameterized with the TIP3P forcefield. Following 10,000 steps of conjugate gradient energy minimization, the system was equilibrated for 12 ns with harmonic restraints on the protein backbone with force constant of 1 kcal/(mol Å^2^) and 50 ns without restraints with a 2 fs time step. The production simulations were run in the NPT ensemble as above. The simulation was run for 600 ns with a GPU-accelerated installation of NAMD v3.0b2 on a GPU node on the SCI cluster at UMass Chan Medical School.

### Analysis of MD simulations

RMSFs, inter-atomic and -residue distances (including for fluorophore centers of mass), and dihedral angles sampled during the simulations of GP^CL^ were calculated using the tcl scripting interface in VMD. Secondary structure was determined using the Timeline plugin in VMD. Residue correlation matrices were calculated using the MDToolbox in Matlab (Matsunaga and Sugita, 2018). The PCA of the simulation trajectory was performed using the ProDy python library (Zhang et al., 2021). For the purposes of PCA, the three replicate simulations of each of the A82 and V82 models were concatenated, as the eigenvectors of the covariance matrices were found to be similar (Aalten et al., 1995). Projection of the simulations into the eigenspace formed by the first three principal components and subsequent clustering using Gaussian mixture models was performed in Matlab. The number of clusters was determined through maximization of likelihood and the Akaike information criterion. All figures were prepared in Matlab and VMD.

### Cell lines and cell culture

HEK-293T (American Type Culture Collection, Manassas, VA) and HEK-293T-FIRB (Mukherjee et al., 2014) cells were maintained in Dulbecco’s Modified Eagle Medium (Gibco, ThermoFisher Scientific, Waltham, MA) supplemented with 10% cosmic calf serum (HyClone, Cytiva Life Sciences, Marlborough, MA), 100 U/mL penicillin, 100 µg/mL streptomycin and 2 mM L-Glutamine (Gibco) at 37°C with 5% CO_2_. The HEK-293T-FIRB cell line was provided by Dr. T. C. Pierson (NIAID, NIH). Expi293F and ExpiCHO-S cells (Gibco) were cultured in Expi293 Expression Medium or ExpiCHO Expression Medium, respectively (Gibco) with orbital shaking at 125 rpm, 37°C with 8% CO_2_.

### Plasmid DNA

For GPΔmuc expression on pseudovirus, the plasmid pGL4.23 WT 2014 EBOV Delta-Mucin-Like-Domain (Addgene plasmid # 86021) was first modified by site-directed mutagenesis (GeneScript USA Inc., Piscataway, NJ) to change the C-terminal *amber* stop codon to an *ochre* stop codon, to prevent readthrough and incorporation of trans-Cyclooct-2-en-L-Lysine axial isomer (TCO*A) at this site during the production of pseudovirus for smFRET imaging. This plasmid was further modified to introduce *amber* stop codons at specific sites for fluorophore attachment, namely N550TAG (asparagine to *amber* stop codon at GPΔmuc amino acid 550) followed by K617TAG (lysine to *amber* stop codon at GPΔmuc amino acid 617) by overlap PCR using *PfuUltra* II Fusion HS DNA Polymerase (Agilent Technologies Inc., SantaClara, CA), the following primers, and the restriction sites AgeI and XbaI:

AgeI-fwd: 5’-ATATCAGGCTACCGGTTTTG-3’
XbaI-rev: 5’-CCGGCCGCCCCGACTCTAGA-3’
N550TAG-fwd: 5’-TAGCAAGATGGTTTAATCTGTGGGTTGA-3’
N550TAG-rev: 5’-CTAGTGCATTAGCCCCTCTG-3’
K617TAG-fwd: 5’-TAGAACATAACAGACAAAATTGATCAGATTATTCATGA-3’
K617TAG-rev: 5’-TCTAGGTCCAATCATGTGGTTCGAT-3’

The A82V mutation (alanine to valine mutation at GPΔmuc amino acid 82) was introduced to the above plasmids by Q5 Site-directed mutagenesis (New England Biolabs, Ipswich, MA) and the primers:

A82V-fwd: 5’-CGTGCCATCTGTGACTAAAAGAT-3’
A82V-rev: 5’-TCAGTTGCCACTCCATTC-3’

For trimeric GPΔTM expression for FCS experiments, the nucleotide sequence encoding GP amino acid residues 1 to 632 from pGL4.23 WT 2014 EBOV Delta-Mucin-Like-Domain was synthesized and cloned by GenScript into the pHL-Sec vector previously described (Durham et al., 2020; Jain et al., 2023), replacing the GPΔTM Mayinga variant sequence. The A1 peptide (GDSLDMLEWSLM) was placed between amino acid 32 and 33 of GP1 and the A4 peptide (DSLDMLEW) placed between 501 and 502 of GP2, analogous to the constructs detailed in (Durham et al., 2020). The A82V mutation was introduced by site-directed mutagenesis (GenScript). Finally, the recognition sequence for the WELQut Protease (ThermoFisher) was added to all GPΔTM expression plasmids by overlap PCR, to specifically remove the glycan cap without non-specific cleavage (and subsequent loss of fluorescence) of the N-terminal A1 and A4 peptides as noted after treatment with the protease Thermolysin.

The GP1 amino acids V203 to T206 (i.e. VNAT) were replaced with the amino acids WELQ using the primers below and the restriction sites EcoRI and AgeI:

EcoRI-fwd: 5’-GCGGCCGTCTCAGGCCGAATTCA-3’
AgeI-rev: 5’-CTCATTAGTTCCAAAACCGGTAGCC-3’
WELQut_fwd: 5’-TGGGAATTGCAAGAGGACCCGTC-3’
WELQut_rev: 5’-TTGCAATTCCCACGGCTCTCTCAAGG-3’

Plasmids for the expression of (i) luminal domain C of NPC1 (Miller et al., 2012) provided by Dr. K. Chandran (Albert Einstein College of Medicine, NY), (ii) NESPylRS^AF^/tRNA^Pyl^ (Nikić et al., 2016) and the dominant negative eRF1 mutant E55D (Schmied et al., 2014) provided by Dr. E. Lemke (Johannes Gutenberg-University of Mainz, Germany) and (iii) HIV-1 GagPol under the control of the cytomegalovirus promotor, provided by Dr. W. Mothes (Yale University School of Medicine, CT) (Das et al., 2020) have been described previously. Expression plasmids for the humanized anti-GP1 H3C8 IgG monoclonal antibody were generated by cloning the variable heavy and light chain fragments into human IgG expression antibody expression vectors, as noted previously (Jain et al., 2023). Specifically, mouse hybridoma cell line expressing the monoclonal antibody M.F88-H3C8 (Ou et al., 2011) was provided by Dr. C. Wilson (FDA, Bethesda, MD). M.F88-H3C8 cells were expanded and provided to GeneScript for identification and synthesis of full-length heavy and light chain sequences (**Table S1**) in the expression vector pcDNA 3.1 (+). Variable regions were then cloned into the human Igγ1 and Igκ expression vectors (Tiller et al., 2008) provided by Dr. M. C. Nussenzweig (The Rockefeller University, NY), which already contained a murine Ig gene signal peptide sequence. The AgeI/SalI or AgeI/BsiWI restriction sites were used for Igγ1 and Igκ expression vector cloning, respectively. All plasmid sequences were confirmed by sequencing (Genewiz, Azenta Life Sciences, South Plainfield, NJ).

### Protein expression & purification

Expression and purification of luminal domain C of NPC1 (sNPC1-C) has been described previously (Vaknin et al., 2024). GPΔTM for FCS experiments was made by transfecting Expi293F cells using polyethyleneimine (PEI MAX, Polysciences, Warrington, PA). The plasmids pHL-Sec-GPΔTM, pHL-Sec-GPΔTM-A1A4 and the expression plasmid for Furin protease were co-transfected at a ratio of 3.33 to 1.66 to 1, to ensure that trimers had a single A1A4-containing protomer on average and contained minimal GP0. At 4 days post-transfection, the cell supernatant was harvested, purified using Ni-NTA Agarose (Invitrogen) using the purification method previously described (Jain et al., 2023). Protein-containing fractions were buffer exchanged/ concentrated in 20 mM HEPES, 50 mM NaCl, pH 7.5 using Vivaspin 6 Centrifugal Concentrators (Sartorius, Gottingen, Germany), then stored at -80°C.

In preparation for FCS experiments, GPΔTM was dye-labelled by overnight incubation at room temperature with 5 μM LD650 conjugated to coenzyme A (LD650-CoA; Lumidyne Technologies, New York, NY), 5 μM acyl carrier protein synthase (AcpS) and 10 mM MgOAc in 20 mM HEPES, 50 mM NaCl, pH 7.5. The glycan cap was then cleaved by incubation with 6 Units of WELQut protease (ThermoFisher) per 1μg of protein at 37°C for 40 minutes, cooled to room temperature and immediately purified by Superdex 200 Increase 10/300 GL column (GE Healthcare, Chicago, IL) size exclusion chromatography in PBS. The peak fraction corresponding to LD650-labelled GP^CL^ΔTM was buffer exchanged/concentrated in 20 mM HEPES, 50 mM NaCl, pH 7.5 using Amicon Ultra Centrifugal Filters (MilliporeSigma, Burlington, MA), aliquoted and stored at -80°C.

The humanized anti-GP1 H3C8 IgG monoclonal antibody was expressed in ExpiCHO-S cells using ExpiFectamine CHO Reagent (ThermoFisher) according to the manufacturer’s standard expression protocol and a 2 to1 ratio of H3C8 Igκ to Igγ1 expression vectors. At 9 days post-transfection, the cell supernatant was harvested and purified using Pierce Protein G Agarose (ThermoFisher) and buffer exchanged/concentrated in PBS to ∼1µg/µl using Vivaspin 6 Centrifugal Concentrators (Sartorius).

### Liposome preparation and fluorescence correlation spectroscopy (FCS)

Methods related to FCS experiments, including liposome preparation and data analysis, were identical to those previously detailed (Jain et al., 2023). Briefly, LD650-labelled GP^CL^ΔTM was incubated with100-nm diameter liposomes at 37°C for 20 min. Liposomes were composed of POPC (1-palmitoyl-2-oleoyl-glycero-3-phosphocholine), POPS (1-palmitoyl-2-oleoyl-sn-glycero-3-phospho-L-serine (sodium salt)), BMP (bis(monooleoylglycero)phosphate (S,R Isomer)) and cholesterol (all from Avanti Polar Lipids, Alabaster, AL). The lipid composition used was POPC:POPS:BMP:Cholesterol (40:40:15:5 mol%) which was shown to robustly support GP-membrane interaction in a pH- and Ca^2+^-dependent manner (Jain et al., 2023). 100 autocorrelation curves of 5 sec each were recorded at room temperature using a 638 nm laser in a CorTector SX100 (LightEdge Technologies Ltd., Zhongshan City, China). Curve fitting and data analysis were previously described (Jain et al., 2023).

### Pseudovirus production

Reporter pseudovirus stocks were produced by transfecting HEK-293T-FIRB cells with Lipofectamine 2000 Transfection Reagent (ThermoFisher). For Western blots and infectivity assays, cells were transfected with a single pGL4.23 GP-expressing plasmid encoding either (i) WT or (ii) 82V GPΔmuc, or with expression plasmids for NESPylRS^AF^/tRNA^Pyl^ and eRF1 E55D in addition to either (iii) WT or (iv) 82V 550/617TAG GPΔmuc. 30 minutes before transfection, the cell culture media was replaced with fresh media, which was also supplemented with 1.2mM TCO*A (SiChem GmbH, Bremen, Germany) for cells transfected with GPΔmuc plasmids containing 550/617TAG. 18 hrs after transfection, cells were transduced with ΔG-VSV-GFP-G, a ΔG-VSV reporter virus expressing GFP (Whitt, 2010) pseudotyped with the VSV Glycoprotein (G protein). After 5 hrs, the cell culture medium was removed, cells were washed with PBS, fresh culture medium was added (with or without TCO*A, as appropriate) and cells were cultured for an additional 18 to 20 hrs. The ΔG-VSV-GFP-GPΔmuc pseudovirus-containing medium was filtered through a 0.45µM pore size filter and concentrated through a 10% sucrose cushion by ultracentrifugation at 25,000 rpm for 2 hrs at 4°C. Viral pellets were resuspended in PBS and stored in aliquots at -80°C.

For smFRET imaging assays, the cell culture media of HEK-293T-FIRB cells was replaced with fresh media supplemented with 1.2mM TCO*A (SiChem GmbH) 30 min before transfection. Cells were co-transfected with 5 plasmids: pGL4.23 GPΔmuc, pGL4.23 GPΔmuc 550/617TAG, NESPylRS^AF^/tRNA^Pyl^, eRF1 E55D and the HIV-1 GagPol expression plasmid. An optimized 1:4 ratio of no TAG to 550/617TAG GPΔmuc plasmids was used to ensure that pseudovirions rarely contained more than a single GPΔmuc 550/617TAG protomer per virion, but samples still had an adequate proportion dye-labelled virus, as the translation efficiency of GPΔmuc 550/617TAG is reduced compared to GPΔmuc. Suppression of the *amber* codons in the GPΔmuc 550/617TAG transcript enabled incorporation of the non-natural amino TCO*A into GPΔmuc. 48 hrs post-transfection, the cell supernatant was harvested, filtered and concentrated through a 10% sucrose cushion as described above. Viral pellets were resuspended in PBS. The glycan cap was proteolytically removed from GPΔmuc by incubation with Thermolysin *Bacillus thermoproteolyticus* (Calbiochem, San Diego, CA) at a final concentration of approximately 0.32 µg/mL for 60 to 65 min at 37°C. Samples were cooled on ice for 10 mins, labelled with 500 nM each of 3-(p-benzylamino)-1,2,4,5-tetrazine-Cy3 and 3-(p-benzylamino)-1,2,4,5-tetrazine-Cy5 (both from Jena Biosciences, Jena, Germany) at room temperature with gentle mixing for 30 mins, before adding 60 μM DSPE-PEG(2000) Biotin (Avanti Polar Lipids) for an additional 30 min. Samples were layered onto a density gradient of 6–30% OptiPrep (Sigma-Aldrich, MilliporeSigma, Burlington, MA) and spun at 35,000 rpm for 1 hr at 4°C. Gradient fractions were collected and analyzed by western blot to confirm the presence of GP1 and complete glycan-cap cleavage. Virus-containing fractions were stored in aliquots at -80°C.

### Western Blots

Pseudovirus samples were heated to 95°C for 5 minutes with LDS Sample Buffer (ThermoFisher) and 10% 2-mercaptoethanol (Fisher Chemical, Hampton, NH), run on a 4-20% precast polyacrylamide gel (Bio-Rad, Hercules, CA) and transferred to a nitrocellulose membrane (Bio-Rad). After blocking overnight at 4°C in 5% skim milk, PBS, 0.1% Tween-20, ThermoFisher) (PBS-T), membranes were incubated for 1 hr at room temperature with anti-GP1 H3C8 HuIgG at a 1:1000 dilution in 5% milk in PBS-T, washed 3 times with PBS-T and incubated for 1 hr at room temperature with horseradish peroxidase-conjugated Goat anti-Human IgG Fc Cross-Adsorbed Secondary Antibody (ThermoFisher) at 1:5000 dilution in 5% milk in PBS-T. Membranes were washed 3 times with PBS-T and developed with SuperSignal West Pico PLUS chemiluminescent substrate (ThermoFisher). Membranes were then stripped for 45 minutes at 60°C in 10% sodium dodecyl sulfate, 0.5M Tris HCl pH6.8, 0.8% 2-mercaptoethanol and blocked in 5% milk in PBS-T as described above. Stripped and blocked membranes were reprobed with the anti-VSV Matrix Protein M Antibody, clone 23H12 (Kerafast, Inc., Shirley MA) followed by horseradish peroxidase-conjugated Rabbit anti-Mouse IgG Fc Secondary Antibody (ThermoFisher) at dilutions of 1:2000 and 1:2500 respectively, in 5% milk in PBS-T and developed as described above. Protein band intensities were quantified with ImageJ software (U.S. National Institutes of Health, Bethesda, MD).

### Infectivity assay

HEK-293T cells were seeded in a 96-well plate at a density of 5.5 × 10^4^ cells/well the day before infection. Concentrated ΔG-VSV-GFP-GPΔmuc pseudovirus was diluted in PBS as necessary and supplemented with 10% cosmic calf serum. The cell culture medium was replaced with virus samples and spun at 300 x *g* for 2 min and cultured for 5.5 hr at 37°C. Cells were harvested by trypsinization, washed with PBS and fixed in 2% paraformaldehyde (ThermoFisher). GFP expression was quantified by flow cytometry using a MACSQuant Analyzer 16 Flow Cytometer (Miltenyi Biotec, Bergisch Gladbach, Germany) and FlowJo v10.10.0 software (BD Life Sciences, Ashland, OR). Two independent infectivity experiments were performed, each using a different pseudovirus preparation. Infectivity was normalized in the following way: densitometric analysis from the Western blot was used to calculate the ratio of EBOV GP1 to VSV M per µl of viral sample loaded. Raw infectivity values were normalized for these differences in GP incorporation levels and expressed relative to wild-type GP infectivity (**Figure S5**).

### smFRET imaging assay and data analysis

Pseudovirions were immobilized on polyethylene glycol (PEG)-passivated, streptavidin-coated quartz slides via DSPE-PEG(2000) Biotin (Avanti Polar Lipids), and imaged using wide-field prism-based TIRF microscopy as previously described (Blakemore et al., 2021; Jain et al., 2023). Imaging was performed at 10 frames/sec using the Micro-Manager microscope control software (micromanager.org) (Edelstein et al., 2014). smFRET trajectories were extracted from movies and analyzed using the SPARTAN software package (Juette et al., 2016). Trajectories that met the following criteria were compiled into histograms and analyzed: FRET was detectable for at least 5 frames prior to photobleaching; the correlation coefficient between donor and acceptor fluorescence traces was less than -0.4; signal-to-noise ratio was greater than 10; and background intensity was less than 50. Trajectories that met these criteria were visually verified and subjected to HMM analysis using the maximum point likelihood (MPL) algorithm (Qin et al., 2000). As indicated in **Figure S2**, the trajectories were fit to multiple models and the model fitness was evaluated by AIC (Akaike, 1974). Through this procedure, a model with five fully connected FRET states was selected; addition of the sixth state did not improve model fitness. The idealization generated by the HMM analysis was used to reconstruct Gaussian distributions, which were overlaid on the FRET histograms, and to calculate the occupancy in each FRET state, which was displayed as a violin plot (**Figure 7**).

Kinetic analysis was performed by extracting the dwell times in the 0.8-FRET state and compiling them into histograms with logarithmically spaced bins to better visualize the multi-modal nature of the distributions (Sigworth and Sine, 1987). The dwell time histograms were fit to the sum of two exponential distributions with amplitudes *A*_slow_ and *A*_fast_, and time constants *t*_slow_ and *t*_fast_, using non-linear least squares fitting. Confidence intervals were calculated from the fit residuals and used to estimate *p*-values for comparisons of the fits. *p*-values less than 0.05 were taken to indicate statistical significance. No statistically significant difference was seen in the time constants across all experimental conditions. Only the amplitudes of the slow and fast components varied.

