## Supplemental Material for "Molecular basis for the increased membrane fusion activity of the Ebola virus glycoprotein A82V variant from the 2013-2016 epidemic: insights from simulations and experiments"

| Heavy Chain | |
|---|---|
| DNA sequence (1407 bp) | **Signal sequence-FR1-CDR1-FR2-CDR2-FR3-CDR3-FR4-Constant region-Stop codon**<br>ATGGAATGGAGCTGGGTCTTTCTCTTCCTCCTGTCAGTAATTGCAGGTGTCCAATCCCAGGTTCAACTGCAGCAGTCTGGGGCTGAACTGGTGAGGCCT<br>GGGGCTTCAGTGACGCTGTCCTGCAAGACTTCGGGCTACACATTTAGTGACTATGAAATACTCTGGGTGAAACAGACACCTGTGCATGGCCTGGAATGG<br>ATTGGAGGTGTTGCTCCTAAAACTGGTGATACTTCCTACAATCAGAACTTCAAGGGCAAGGCCATATTGACTGCAGACAAGTCCTCCAGAGCAGCCTAC<br>ATGGAACTCCGCCGCCTGACATCTGAGGACTCTGCCGTCTATTACTGTACAACCGAGGTCTACTCTAGGTACGACGGAGACCCTTACTACGCTATGGAC<br>TATTGGGGTCAAGGAACCGCAGTCACCGTCTCCTCAGCCAAAACGACACCCCCATCTGTCTATCCACTGGCCCCTGGATCTGCTGCCCAAACTAACTCC<br>ATGGTGACCCTGGGATGCCTGGTCAAGGGCTATTTCCCTGAGCCAGTGACAGTGACCTGGAACTCTGGATCCCTGTCCAGCGGTGTGCACACCTTCCCA<br>GCTGTCCTGCAGTCTGACCTCTACACTCTGAGCAGCTCAGTGACTGTCCCCTCCAGCACCTGGCCCAGCGAGACCGTCACCTGCAACGTTGCCCACCCG<br>GCCAGCAGCACCAAGGTGGACAAGAAAATTGTGCCCAGGGATTGTGGTTGTAAGCCTTGCATATGTACAGTCCCAGAAGTATCATCTGTCTTCATCTTC<br>CCCCCAAAGCCCAAGGATGTGCTCACCATTACTCTGACTCCTAAGGTCACGTGTGTTGTGGTAGACATCAGCAAGGATGATCCCGAGGTCCAGTTCAGC<br>TGGTTTGTAGATGATGTGGAGGTGCACACAGCTCAGACGCAACCCCGGGAGGAGCAGTTCAACAGCACTTTCCGCTCAGTCAGTGAACTTCCCATCATG<br>CACCAGGACTGGCTCAATGGCAAGGAGTTCAAATGCAGGGTCAACAGTGCAGCTTTCCCTGCCCCCATCGAGAAAACCATCTCCAAAACCAAAGGCAGA<br>CCGAAGGCTCCACAGGTGTACACCATTCCACCTCCCAAGGAGCAGATGGCCAAGGATAAAGTCAGTCTGACCTGCATGATAACAGACTTCTTCCCTGAA<br>GACATTACTGTGGAGTGGCAGTGGAATGGGCAGCCAGCGGAGAACTACAAGAACACTCAGCCCATCATGGACACAGATGGCTCTTACTTCGTCTACAGC<br>AAGCTCAATGTGCAGAAGAGCAACTGGGAGGCAGGAAATACTTTCACCTGCTCTGTGTTACATGAGGGCCTGCACAACCACCATACTGAGAAGAGCCTC<br>TCCCACTCTCCTGGTAAATGA |
| Amino acid sequence (468 aa) | **Signal peptide-FR1-CDR1-FR2-CDR2-FR3-CDR3-FR4-Constant region-Stop codon**<br>MEWSWVFLFLLSVIAGVQSQVQLQQSGAELVRPGASVTLSCKTSGYTFSDYEILWVKQTPVHGLEWIGGVAPKTGDTSYNQNFKGKAILTADKSSRAAY<br>MELRRLTSEDSAVYYCTTEVYSRYDGDPYYAMDYWGQGTAVTVSSAKTTPPSVYPLAPGSAAQTNSMVTLGCLVKGYFPEPVTVTWNSGSLSSGVHTFP<br>AVLQSDLYTLSSSVTVPSSTWPSETVTCNVAHPASSTKVDKKIVPRDCGCKPCICTVPEVSSVFIFPPKPKDVLTITLTPKVTCVVVDISKDDPEVQFS<br>WFVDDVEVHTAQTQPREEQFNSTFRSVSELPIMHQDWLNGKEFKCRVNSAAFPAPIEKTISKTKGRPKAPQVYTIPPPKEQMAKDKVSLTCMITDFFPE<br>DITVEWQWNGQPAENYKNTQPIMDTDGSYFVYSKLNVQKSNWEAGNTFTCSVLHEGLHNHHTEKSLSHSPGK- |
| **Light Chain** | |
| DNA sequence (717 bp) | **Signal sequence-FR1-CDR1-FR2-CDR2-FR3-CDR3-FR4-Constant region-Stop codon**<br>ATGAGTCCTGCCCAGTTCCTGTTTCTGTTAGTGCTCTGGATTCGGGAAACCAACGGTGATGTTGTGATGACCCAGACTCCACTCACTTTGTCGGTTACC<br>ATCGGACAACCAGCCTCCATCTCTTGCAAGTCAAGTCAGAGCCTCTTAGATAGTGATGGAAGGACATATTTGAATTGGTTGTTACAGAGTCCAGGCCAG<br>TCTCCGAAGCGCCTAATATATCTGGTGTCTAGACTGGACTCTGGAGTCCCTGACAGGTTCACTGGCAGTGGATCAGGGACAGATTTCACACTGAAAATC<br>AGCAGAGTGGCGGCTGAGGATTTGGGAGTTTATTATTGCTGGCAAGGTACACATTTTCCTCAGACGTTCGGTGGAGGCACCAAGCTGGAAATCAAACGG<br>GCTGATGCTGCACCAACTGTATCCATCTTCCCACCATCCAGTGAGCAGTTAACATCTGGAGGTGCCTCAGTCGTGTGCTTCTTGAACAACTTCTACCCC<br>AAAGACATCAATGTCAAGTGGAAGATTGATGGCAGTGAACGACAAAATGGCGTCCTGAACAGTTGGACTGATCAGGACAGCAAAGACAGCACCTACAGC<br>ATGAGCAGCACCCTCACGTTGACCAAGGACGAGTATGAACGACATAACAGCTATACCTGTGAGGCCACTCACAAGACATCAACTTCACCCATTGTCAAG<br>AGCTTCAACAGGAATGAGTGTTAG |
| Amino acid sequence (238 aa) | **Signal peptide-FR1-CDR1-FR2-CDR2-FR3-CDR3-FR4-Constant region-Stop codon**<br>MSPAQFLFLLVLWIRETNGDVVMTQTPLTLSVTIGQPASISCKSSQSLLDSDGRTYLNWLLQSPGQSPKRLIYLVSRLDSGVPDRFTGSGSGTDFTLKI<br>SRVAAEDLGVYYCWQGTHFPQTFGGGTKLEIKRADAAPTVSIFPPSSEQLTSGGASVVCFLNNFYPKDINVKWKIDGSERQNGVLNSWTDQDSKDSTYS<br>MSSTLTLTKDEYERHNSYTCEATHKTSTSPIVKSFNRNEC- |

**Table S1. Consensus full-length antibody sequences of M.F88-H3C8 mouse hybridoma cells.** Total RNA was isolated from the hybridoma cells following the technical manual of TRIzol® Reagent. Total RNA was then reverse-transcribed into cDNA using either isotype-specific anti-sense primers or universal primers following the technical manual of PrimeScript™ 1st Strand cDNA Synthesis Kit. Antibody fragments of heavy chain and light chain were amplified according to the standard operating procedure (SOP) of rapid amplification of cDNA ends (RACE) of GenScript. Amplified antibody fragments were cloned into a standard cloning vector separately. Colony PCR was performed to screen for clones with inserts of correct sizes.

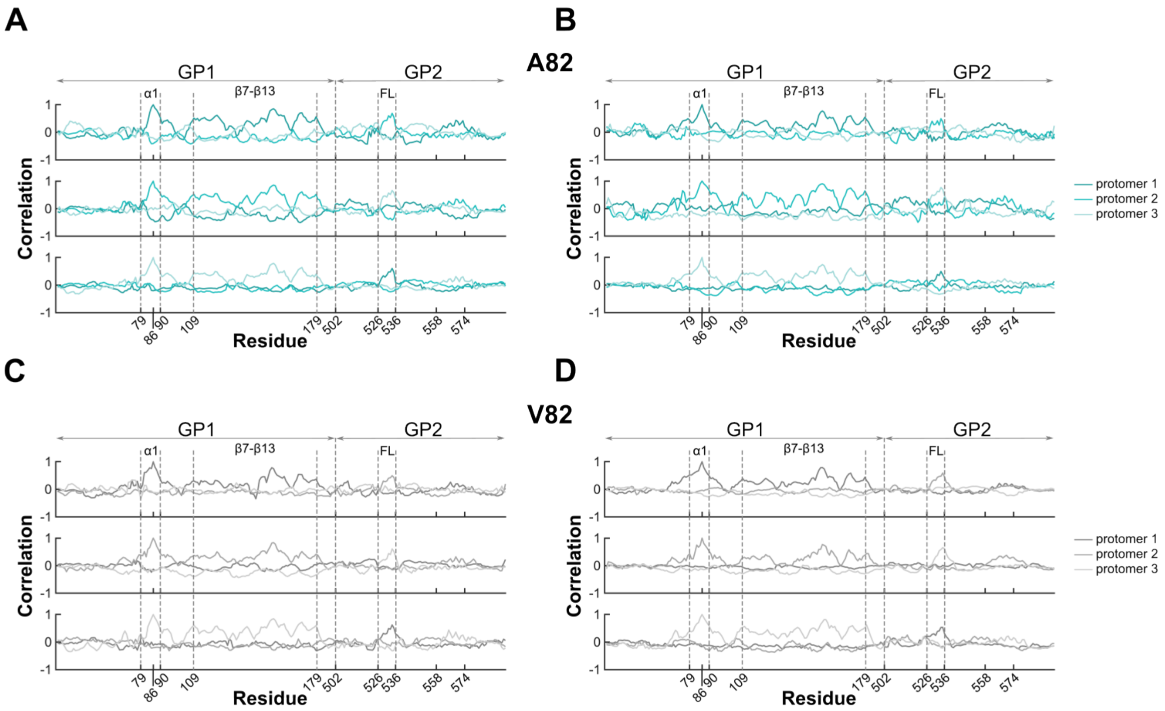


**Figure S1. Correlation of W86 and GP$^{CL}$ fusion loop dynamics.** (**A**) Correlation coefficients (cc) of W86 with all residues within each of the three protomers, determined from one replicate MD simulation of A82 GP$^{CL}$. Key residues and structural features are indicated. High correlation coefficient is seen between W86 and the $\alpha$1 helix (cc > 0.7) and $\beta$7-$\beta$13 sheets (including H154 in $\beta$10; cc $\approx$ 0.8) in the same protomer, and the fusion loop (FL) of a neighboring protomer (cc $\approx$ 0.6). (**B**) The same data from the third replicate A82 simulation. (**C-D**) The same data from two replicate simulations of the V82 GP$^{CL}$ trimer. Equivalently high correlation was seen between W86 and the $\alpha$1 helix, $\beta$7-$\beta$13 sheets, and the FL as for the A82 simulations.

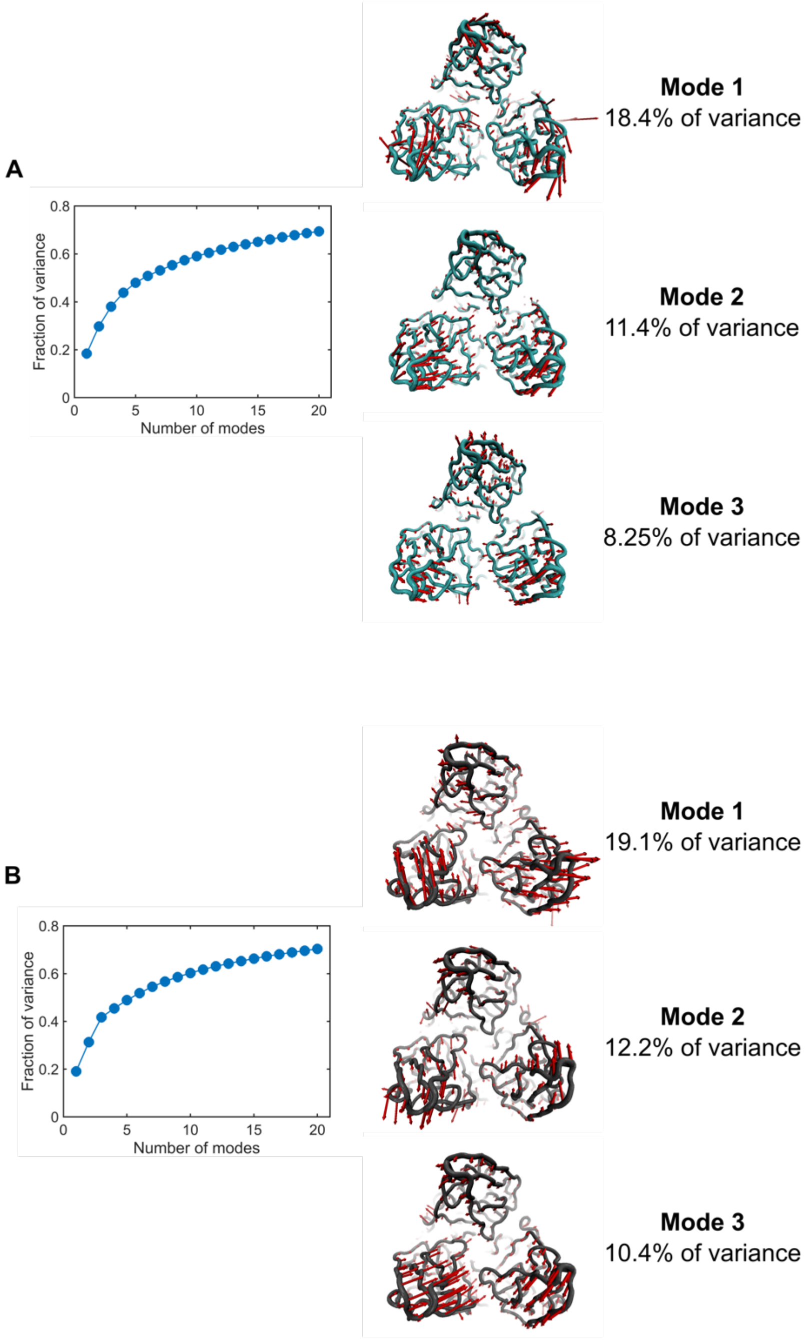


**Figure S2.** (**A**) The fraction of variance attributed to each of the modes determined through PCA of the MD simulation of the A82 $GP^{CL}$ trimer. At right are structural depictions of the modes that account for the indicated fractional variance. Approximately 38% of the variance is accounted for by the first three modes. (**B**) The same analysis of the V82 $GP^{CL}$ simulation. Approximately 42% of the variance is accounted for by the first three modes.

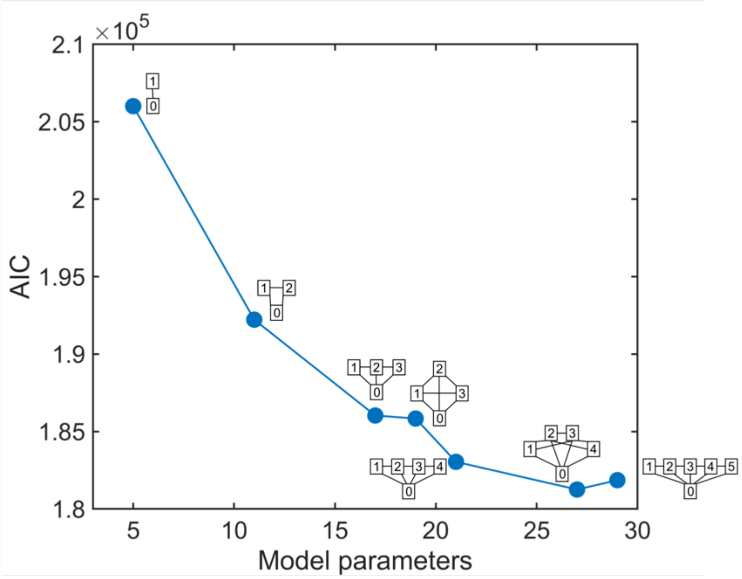


**Figure S3. Model selection through minimization of the AIC.** smFRET trajectories from A82 GP$^{CL}$ were fit to a series of different models by maximum likelihood optimization using the MPL algorithm (see Materials and Methods). The maximized likelihood estimated for each model was corrected for differing numbers of model parameters using the AIC. Under this procedure, the model with best fitness among those considered will generate minimum AIC values. This criterium identified a 5-state fully connected model as providing the best representation of the data among the models considered; no improvement in model fitness was seen upon addition of a sixth state. Overlaid on the plot are schematic representations of the kinetics models considered. In all models, the 0 state corresponds to the 0-FRET state, with the others reflecting non-zero FRET states. Lines represent connections (allowed transitions) between states. All non-zero FRET states are connected to the 0-FRET state since photobleaching is seen from all FRET states.

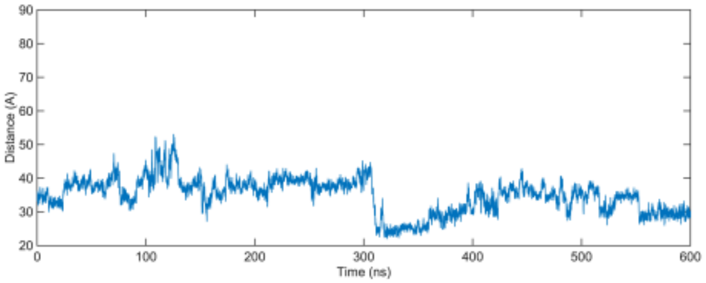


**Figure S4. The distance between Cy3 and Cy5 bound to GP$^{CL}$.** An atomic molecular model of GP$^{CL}$ (PDB: 5JQ3) with fluorophores attached to positions 550 and 617 was simulated for 600 ns (see Materials and Methods). The distance between the center of mass of the fluorophores was calculated for each frame of the simulation, yielding an average distance of 36±5 Å.

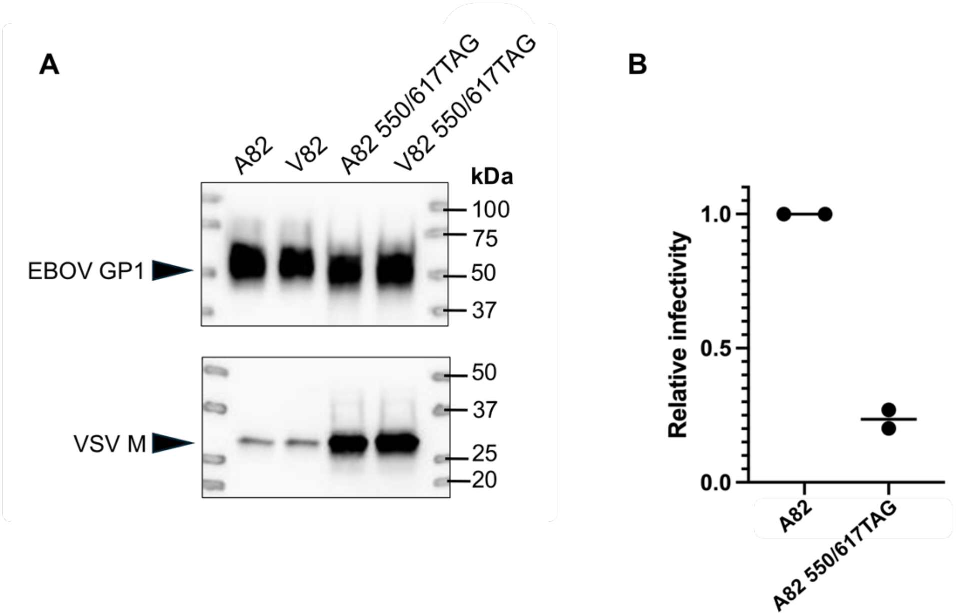


**Figure S5. Evaluation of GP with amber codons at positions 550 and 617.** (**A**) Western blots probing for EBOV GP1 and VSV M. (**B**) Infectivity of VSV pseudovirions containing only wild-type A82 GP, or only modified A82 GP with amber codons at 550 and 617. Each point indicates the arithmetic mean of two technical replicates. The average of two independent experiments is displayed, normalized to values for wild-type A82 GP as described in Materials and Methods.
